# Remote sensing-based forest modeling reveals positive effects of functional diversity on productivity at local spatial scale

**DOI:** 10.1101/2022.08.11.503677

**Authors:** Fabian D Schneider, Marcos Longo, Eugénie Paul-Limoges, Victoria M Scholl, Bernhard Schmid, Felix Morsdorf, Ryan P Pavlick, David S Schimel, Michael E Schaepman, Paul R Moorcroft

**Affiliations:** Jet Propulsion Laboratory, California Institute of Technology, Pasadena, CA 91109, United States; Remote Sensing Laboratories, Department of Geography, University of Zurich, CH-8057 Zurich, Switzerland; Lawrence Berkeley National Laboratory, Climate and Ecosystem Sciences Division, Berkeley, CA, 94720, United States; Earth Lab, Cooperative Institute for Research in Environmental Sciences, University of Colorado Boulder, Boulder, CO 80303, United States; Department of Geography, University of Colorado Boulder, Boulder, CO 80309, United States; Department of Organismic and Evolutionary Biology, Harvard University, Cambridge, MA 02138, United States

**Keywords:** biodiversity, ecosystem functioning, temperate forest, imaging spectroscopy, lidar, terrestrial biosphere modeling, plant functional traits, functional diversity

## Abstract

- Forest biodiversity is critical for many ecosystem functions and services at plot scale, but it is uncertain how biodiversity influences ecosystem functioning across environmental gradients and contiguous larger areas. We used remote sensing and process-based terrestrial biosphere modeling to explore functional diversity–productivity relationships at multiple scales for a heterogeneous forest site in Switzerland.
- We ran the biosphere model with empirical data about forest structure and composition derived from ground-based surveys, airborne laser scanning and imaging spectroscopy for the years 2006–2015 at 10*×*10-m spatial resolution. We then related the model outputs forest productivity to functional diversity under observed and experimental model conditions.
- Functional diversity increased productivity significantly (p *<* 0.001) across all simulations at 20*×*20-m to 30*×*30-m scale, but at 100*×*100-m scale positive relationships disappeared under homogeneous soil conditions.
- Whereas local functional diversity was an important driver of productivity, environmental context (especially soil depth, texture and water availability) underpinned the variation of productivity (and functional diversity) at larger spatial scales. Integration of remotely-sensed information on canopy composition and structure into terrestrial biosphere models helps fill the knowledge gap about how plant biodiversity affects carbon cycling and biosphere feedbacks onto climate over large contiguous areas.

## Introduction

Biodiversity is a key property of forests that affects important ecosystem services, including provisioning services such as timber supply or water purification, cultural services of recreational and spiritual value, and regulating services such as carbon sequestration (Hooper *et al*., 2005; Cardinale *et al*., 2012; Chamagne *et al*., 2017; Aerts *et al*., 2018; Isbell *et al*., 2017). Biodiversity also plays an important role in the carbon cycle and influences vegetation–atmosphere interactions and feedbacks (Schimel *et al*., 2015). For example, higher biodiversity may increase the resistance and resilience of ecosystems to climate change and increase carbon sequestration contributing to climate change mitigation (Isbell *et al*., 2015; Liang *et al*., 2016; Liu *et al*., 2018; Huang *et al*., 2018). Conversely, losses in biodiversity may contribute to accelerating impacts of climate change on forest ecosystems (Mori, 2020), triggering potentially additional species losses (Trisos *et al*., 2020). Despite this recognized importance of plant diversity to help forest ecosystems provide multiple ecosystem functions and services (Díaz *et al*., 2019), there is still considerable uncertainty in the mechanistic relationship between biodiversity and ecosystem functioning (BEF); specifically, (1) may scale up from plot-level to large contiguous areas of forest, (2) how BEF may be modified by environmental variation and change, (3) which measures of biodiversity should be used for large-scale assessments, and (4) how biodiversity effects are represented in demographic modeling of forest dynamics.

Regarding (1), positive effects of biodiversity on ecosystem functioning and stability have been reported widely at plot scale, initially mainly from experimental studies (Balvanera *et al*., 2006; Cardinale *et al*., 2011; Tilman *et al*., 2014) but recently also from observational studies (e.g. Grace *et al*., 2016; Liang *et al*., 2016; Duffy *et al*., 2017). Even though these plot-scale studies may cover areas up to global extent (Liang *et al*., 2016), they are still considering separate plots of particular size and very few studies explicitly test plot-size effects (e.g. Huang *et al*., 2018; Chisholm *et al*., 2013; Poorter *et al*., 2015). Scaling up BEF relationships remains a major challenge (Gonzalez *et al*., 2020). Regarding (2), there have been a number of studies addressing environmental context in BEF studies (see Hong *et al*., 2022, for a recent meta-analysis), but few focused on forests. Paquette & Messier (2011), Ratcliffe *et al*. (2017), and Mina *et al*. (2017) found that the mechanisms and BEF effects varied with environmental conditions and stress, but large uncertainty remains with regard to climate change – even whether forests will remain a carbon sink is uncertain (Brienen *et al*., 2015; Sabatini *et al*., 2019; Arora *et al*., 2020). For (3), previous BEF literature explored which components of biodiversity are most predictive for ecosystem functioning, but no single component is consistently the best (Craven *et al*., 2018). Functional diversity is often considered promising (Loreau, 2000; Hooper *et al*., 2005; Flynn *et al*., 2011), yet rarely assessed explicitly and independent of taxonomy (e.g. Schneider *et al*., 2017; Guilĺen-Escribà *et al*., 2021). Regarding mechanisms, the theoretical reasoning suggests that niche complementarity (complementary traits lead to better resource use of a community as a whole), selection probability (higher likelihood to have highly productive individuals), and ecological insurance (higher likelihood to adapt to or recover from fluctuating environmental conditions or disturbance) underpin biodiversity effects on ecosystem functioning and stability (Tilman *et al*., 1997; Yachi & Loreau, 1999; Loreau, 2000). However, regarding (4), integration of observed biodiversity patterns into terrestrial biosphere models and direct demographic modeling of these mechanisms still represents a challenge, but it is critically important.

Simulations and model experiments using process-based models can extend our knowledge of diversity–productivity relationships and their role in the global carbon cycle. Morin *et al*. (2011), for example, used a process-based dynamic vegetation model to simulate biodiversity experiments for a range of sites and species in Europe, finding consistent positive effects of biodiversity on productivity due to complementarity. Extending this approach to larger scales, Levine *et al*. (2016) and Longo *et al*. (2018) found that ecosystem heterogeneity and diversity were important to predict forest resilience to climate change including extreme drought events in the Amazon. Many terrestrial biosphere models, however, are not able to resolve this level of detail, or only allow to simulate potential vegetation, which can lead to uncertainties in the prediction of the carbon cycle (Braghiere *et al*., 2019) and climate change feedbacks (Schimel *et al*., 2019; Fisher & Koven, 2020). The integration of remote observations, such as from imaging spectroscopy and lidar, of forest composition and structure into models holds great promise to improve our understanding of patterns and drivers of forest diversity and productivity. Previous studies have shown that incorporating such measurements into biosphere models substantially improved their predictions of forest carbon fluxes (Antonarakis *et al*., 2014, 2022).

In this study, we use high-spatial resolution airborne laser scanning and imaging spectroscopy measurements of temperate forest composition and structure to produce initial conditions for the process-based terrestrial biosphere model ED2 (Longo *et al*., 2019b). We then use the remote-sensing initialized model to investigate the role of morphological and physiological diversity, derived from trait outputs by the model after integration of the remote sensing data, on spatial patterns of forest productivity at local to landscape scales (Fig. 1). We use this approach to explore what is driving spatial variation in forest gross primary productivity under observed climatic conditions from 2006 to 2015. Analyses of the model outputs enabled us to address these two main research questions: (1) What is the relationship between morphological and physiological diversity and gross primary productivity (GPP) at different spatial scales? (2) How does the relationship between diversity and productivity vary across environmental and ecological gradients involving different soil types, plant functional and structural types, and vegetation dynamics? To address these questions, we first evaluate the model’s ability to simulate carbon dynamics and seasonality of the studied temperate mixed forest as measured by a fluxtower at the Laegern forest in Switzerland. Second, we run realistic simulations and model experiments to study diversity–productivity relationships across 15 forest sites (Fig. 2). Finally, we discuss current limitations and ways to improve the representation of plant functional diversity in process-based models.

**Fig. 1:**
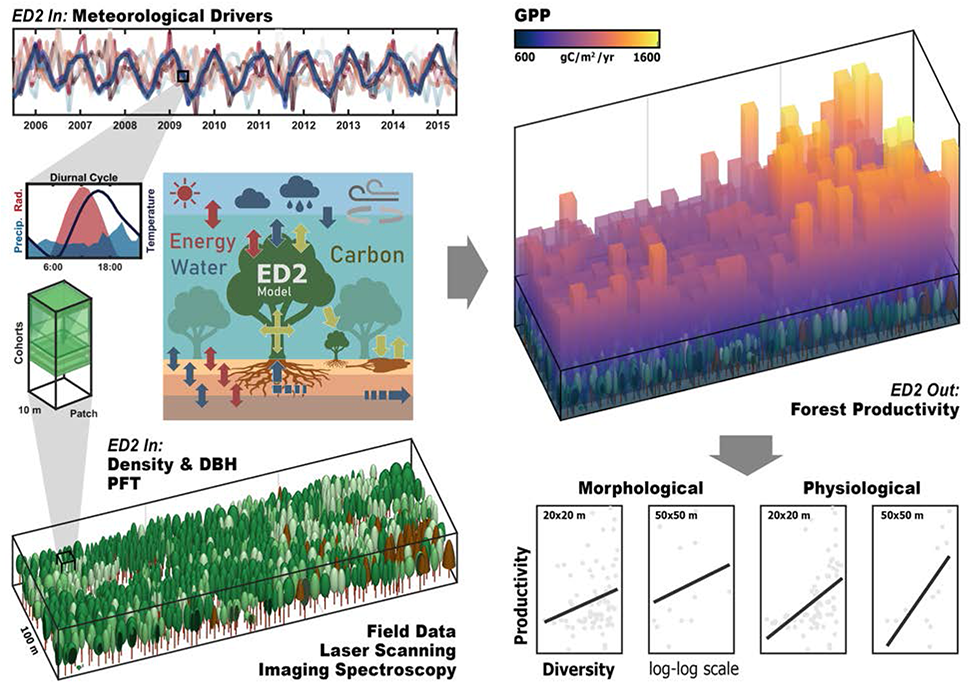
Forest modeling approach to derive forest diversity and productivity relationships in a temperate mixed forest. We used airborne imaging spectroscopy and light detection and ranging (LiDAR) together with field inventory data to derive plant functional types (PFT), tree diameters at breast height (DBH) and stem density. These variables were used to represent the forest in the ED2 terrestrial biosphere model in 10*×*10 m patches and vertically resolved cohorts within patches. Forest productivity was then simulated as gross primary productivity (GPP) based on hourly meteorological drivers and averaged over 10 years, allowing us to relate the underlying morphological and physiological diversity to the simulated long-term forest productivity. The visualization shows an example based on data and simulations conducted in this study.

**Fig. 2:**
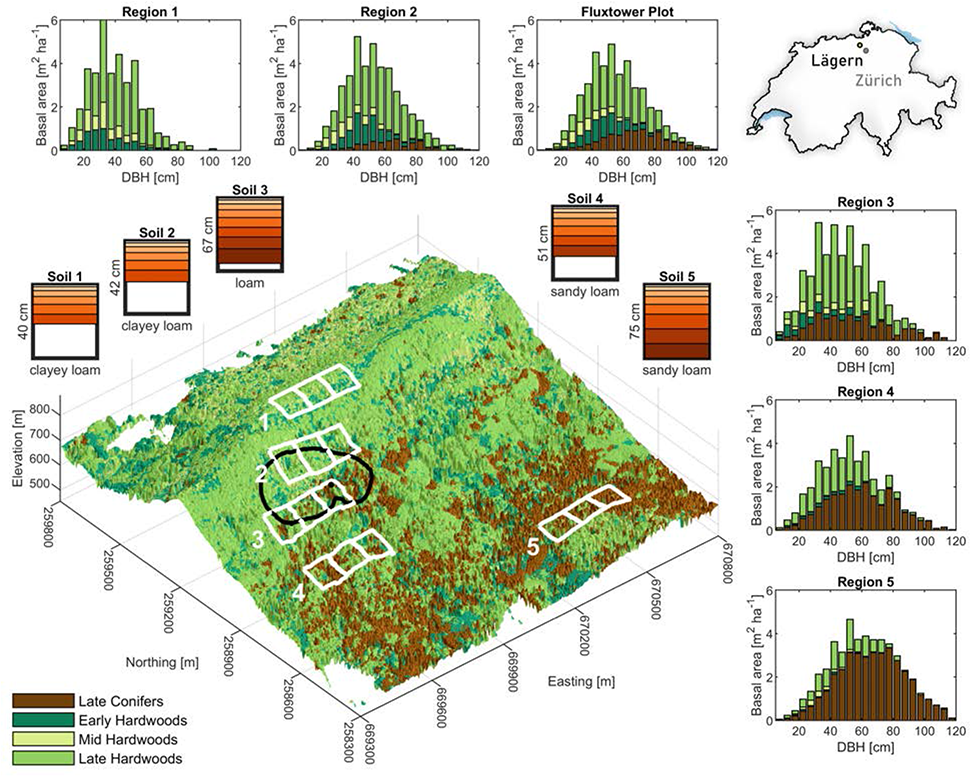
Study regions at the Laegern mountain, Switzerland, covering a gradient in soil, structure, composition and diversity of the temperate mixed forest. The three-dimensional surface topography is colored by the four plant functional types: late-successional conifers, early-, mid- and late-successional hardwoods. Regions 1–5 are each comprised of three 1-ha sites (white boxes) and show a shift from shorter deciduous broadleaf forest on top of the mountain (1) to taller evergreen coniferous forest at lower elevation (5) following a gradient in soils from shallow clayey loam (1) to deeper sandy loam (5). The histograms show the distribution of basal area per tree diameter at breast height (DBH) for the four plant functional types. The Laegern fluxtower is located at the north-east corner of region three (black dot). Model evaluation was performed using forest composition and structure representative of the fluxtower footprint, shown as black outline at 70% flux density. The map (top-right) shows the location of Laegern north-west of Zurich, with respect to the country border of Switzerland.

## Materials and methods

In the following sections, we provide a detailed description and methodology of the study area, the terrestrial biosphere model, the integration of remote sensing data into the model, model evaluation and simulations, and how we derived and analyzed the model outputs forest functional traits, diversity and productivity. In short, we first evaluated the model performance at a fluxtower site in the center of the study area (Fig. 2), which built the basis for the subsequent diversity–productivity analyses. For these analyses, we ran 10-year simulations of forest dynamics and productivity across five regions with varying soil properties in simulations where remote observations were used to initialize the model, and in model experiments using uniform soils (mono-soil), forest composition (mono-cultures) and structure (mono-structure). We calculated traits at the forest patch-level from the model at 10*×*10-m spatial resolution, i.e. (1) basal area, leaf area index and foliage height diversity (*morphological traits*), and (2) specific leaf area (SLA) and carboxylation rate (VC*_max_*) (*physiological traits*) from model-PFT mixtures. We then derived morphological and physiological diversity (richness, divergence and evenness following Schneider *et al*. (2017)) at spatial extents of 20*×*20-m to 100*×*100-m and related them to simulated annual GPP averaged over 2006–2015. Finally, we analyzed the simulated diversity–productivity relationships using general linear models and random forest models.

### Study area

We conducted this study at the Laegern temperate mixed forest in Switzerland (47*^◦^*28’43.0 N, 8*^◦^*21’53.2 E), which is located on a mountain range representing broad ecological and environmental gradients (Fig. 2). The study area is characterized by increasing soil depth (40 to 75 cm) and soil grain size (clayey to sandy loam), and increasing mean tree stem diameter and abundance of conifers from high to low elevation (850–450 m above sea level). The vegetation consists of diverse beech forests with a total of 13 tree species. The dominant deciduous species are common beech (*Fagus sylvatica* L.), European ash (*Fraxinus excelsior* L.), and sycamore maple (*Acer pseudoplatanus* L.). The dominant coniferous species are Norway spruce (*Picea abies* (L.) H. Karst) and silver fir (*Abies alba* Mill.).

The study area has a fluxtower (CH-LAE; Paul-Limoges *et al*., 2020) equipped with eddy-covariance instruments and a meteorological station, as well as a 5.5-ha field plot, where 1307 canopy trees were mapped by stem location, crown extent, diameter at breast height (DBH) and species identity (Morsdorf *et al*., 2020; Guillén-Escribà *et al*., 2021). We selected five regions to study diversity–productivity relationships, covering the range of soil characteristics and the ecological gradient from shorter deciduous broadleaf forest on top of the ridge to taller mixed communities in lower elevations. Each region was sub-divided into three 100*×*100-m sites, selected to minimize environmental heterogeneity within regions and maximize the range of abiotic and biotic variation between regions and sites, respectively (Fig. 2).

### Model description

The ED2 model is a process-based terrestrial biosphere model that represents individual plant-level dynamics (growth, mortality and recruitment), and associated ecosystem-level carbon, water and energy fluxes over time scales ranging from hours to centuries (Fig.1; Longo *et al*., 2019b). In contrast to conventional ‘ecosystem as big leaf’ models that represent the canopy in a highly aggregated manner, the ecosystem demography (ED) model utilizes the concept introduced by Moorcroft *et al*. (2001), in which vegetation is represented as cohorts of plants of similar height and plant functional type, grouped in patches that can be initialized with spatially explicit remote observations, within a site of homogeneous environmental conditions with regard to soil, topography and meteorology (Longo *et al*., 2019b). Plant growth, mortality and recruitment dynamics are simulated as part of vegetation dynamics. The model can be run with vegetation dynamics off, where only phenological and physiological dynamics are simulated. The size- and age-structured representation allows us to incorporate fine-scale spatial variation in canopy structure and composition arising from disturbance processes and spatially-localized, size-dependent competition for light within the above-ground plant canopy and below-ground for water (bigger trees have deeper roots and thus can access water in deeper soil layers). To improve vegetation dynamics regarding canopy structure and light availability compared to previous ED modeling schemes, heterogeneity in horizontal and vertical micro-climate environments has been introduced in ED2 (Medvigy *et al*., 2009; Longo *et al*., 2019b). The ED2 biosphere model has shown good conservation of energy, water and carbon (Longo *et al*., 2019b,a) and has been successfully run with initial conditions from forest-stand data as observed from remote sensing (Longo *et al*., 2020), field plots (Meunier *et al*., 2021) or a combination of both (Bogan *et al*., 2019), although at coarser spatial scales than presented here.

### Integration of remote sensing data into the model

We derived forest composition (PFTs) and structure (DBH and stem density) using field data with remote observations, i.e. airborne imaging spectroscopy and laser scanning (ALS), at high spatial resolution (2*×*2-m), and integrated these empirical data into the model as initial conditions of tree cohorts within forest patches (10*×*10-m).

**Forest composition** was integrated into the model based on a classification of ED2-PFTs originally defined for Harvard forest (Medvigy *et al*., 2009), namely: late-successional conifers and early-, mid-, and late-successional hardwoods (LCf, EHw, MHw, LHw, respectively). These four ED2-PFTs and corresponding physiological traits have been extensively used and tested in temperate forests in North America and Europe (e.g. Medvigy *et al*., 2009, 2010; Antonarakis *et al*., 2014; Jin *et al*., 2017; Paul-Limoges *et al*., 2020). We classified PFTs at 2*×*2-m resolution using a random forest classifier with remotely sensed input features, trained and validated based on field data of 1307 identified trees (Morsdorf *et al*., 2020) and 73 forest stand polygons (Schneider *et al*., 2017). From 81 possible input features, we selected 27 relevant features that yielded the best model results, including morphological and physiological forest traits (Schneider *et al*., 2017), principal components of surface reflectance and continuum-removed reflectance acquired by the airborne imaging spectrometer APEX (Schaepman *et al*., 2015). The classifier had an overall accuracy of 74% and a Cohen’s kappa coefficient of 61% for the classification of plant functional types over the whole forest. Further details on the classification can be found in Supplementary Notes 1 and 2, and Figures S1, S3 and S4.

**Forest structure** was integrated into the model as initial conditions of DBH and stem density estimated from ALS and field data. DBH values were estimated from ALS-derived canopy height using an exponential model that was fitted to field inventory data of 159 late conifers, 253 early hardwoods, 328 mid hardwoods and 566 late hardwoods:

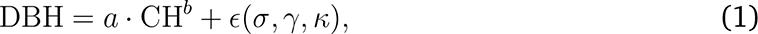

where CH is canopy height, *a* and *b* are coefficients and *ɛ* is the residual distribution around the mean. To predict not only the mean but a realistic DBH distribution, we derived the standard deviation *σ*, skewness *γ* and kurtosis *κ* of the distribution to model *ɛ* (Matlab R2017a, pearsrnd). The resulting values for *a*, *b*, *σ*, *γ* and *κ* are summarized in Table S1. We applied this approach to predict a DBH value for each 2*×*2-m pixel based on its canopy height. Single-tree delineation was applied to the ALS point cloud to estimate the density of stems of different sizes (i.e. canopy height classes) across each 1-ha site. The method had a detection rate of 79% overall, 83% for trees *≥* 35 m and 76% for trees *<* 35 m. We assigned stem density to each 2*×*2-m pixel based on canopy height, by assigning the average density estimate per 1-m canopy height class derived at each study region.

Finally, values were aggregated as cohorts in discrete 10*×*10-m forest patches across each 100*×*100-m site by averaging DBH and stem density for each plant functional type and 10-cm DBH class. This approach reduces the number of cohorts to simulate in the model, reduces computational expenses, and better approximates the individual tree scale of the forest while preserving plant functional type distributions within patches. For more detail see Supplementary Note 3, Figures S1, S2, and predicted stem density and count (Fig. S5) compared with field data (Fig. S6).

**Meteorological drivers** from hourly meteorological variables measured between 2006 and 2015 were used as boundary conditions. Atmospheric pressure at field elevation (hPa), air temperature (*^◦^*C), relative humidity (%), average wind speed (m s*^−^*^1^), and precipitation (mm) were measured at 10-minute intervals on the Laegern fluxtower as part of the national air pollution monitoring network (NABEL) of Switzerland. Incoming shortwave (W m*^−^*^2^) and longwave radiation fluxes (W m*^−^*^2^) were measured at 1-minute intervals. For gap filling of missing values (*<*2%), we used meteorological data from nearby meteorological stations (Supplementary Note 4, Figure S7). All meteorological variables were averaged to one hour for input into the ED2 model. Atmospheric CO_2_ was assumed constant at 391 *µ*mol mol*^−^*^1^, consistent with the average background values observed at Mauna Loa between 2006 and 2015 (Tans & Keeling, 2020).

**Soil** information was used from FOAG (1996) to specify spatial variation in soil depth and texture (relative sand, loam, clay content) as the main ED2 inputs (Fig. 2).

### Model evaluation

We evaluated the model’s carbon flux predictions for the period 2006–2015 against corresponding estimates from eddy-covariance fluxes (Baldocchi, 2003) measured continuously at the Laegern fluxtower. Flux quality post-processing was done following Vickers & Mahrt (1997). Standardized gap filling and partitioning of the net ecosystem CO_2_ exchange into GPP and ecosystem respiration were done using the method from Barr *et al*. (2004). We ran the flux footprint prediction model FFP (Kljun *et al*., 2015) to estimate the average proportions of PFTs, DBH and stem density within the footprint based on the 2*×*2-m remote sensing maps (Figs. S1, S8, S9). We then selected a sample of 200 representative forest patches for simulation in ED2 and comparison to fluxtower estimates. We ran a 10-year spin-up phase using the 200 patches as initial condition before simulating the years 2006–2015 including vegetation dynamics.

We also assessed the predicted phenology by comparing ED2’s monthly simulated LAI with satellite-based LAI estimated using the data assimilation model PhenoAnalysis (Stöckli *et al*., 2011). We ran the PhenoAnalysis model in the data assimilation mode to fit an LAI time series to the noisy Moderate Resolution Imaging Spectroradiometer (MODIS) satellite data taking into account the gap filled meteorological variables from the fluxtower (Fig. S10).

### Model simulations and experiments

For all simulations, we ran the model over a 20-year period by cycling through the meteorological driver time series twice. The first 10-year cycle was considered spin-up, so that the soil moisture and the vegetation dynamics could stabilize after integrating initial conditions of the forest, and only the second 10-year cycle was considered in the analyses.

To address the two research questions and disentangle the effects of forest structure, composition and soil on productivity, we ran five sets of simulations on the 15 (5*×*3) 100*×*100-m sites, summarized in Table 1. The first set of realistic simulations were run with the observed composition, structure, and soil information for the 15 sites, with both vegetation dynamics on and off. We then ran three experimental simulations to which we refer as mono-soils, monocultures and mono-structures. In the mono-soil simulations, the 15 sites were run with observed stand composition and structures on nine uniform soils (3 soil depths *×* 3 soil textures). In the monoculture simulations, the 15 sites were run with four different uniform PFTs, each on 3 soils (i.e. 4 PFTs *×* 3 soils *×* 15 sites). Finally, the mono-structure simulations were run with the observed composition across the 15 sites, but using three different uniform canopy structures, each on 3 soils (3 structures *×* 3 soils *×* 15 sites). Canopy structures were created by replicating the vegetation structure of forest patches that had low (19 m^2^ ha*^−^*^1^), medium (39 m^2^ ha*^−^*^1^) and high (83 m^2^ ha*^−^*^1^) basal area across all 15 sites, while keeping the PFT-composition as observed. The model experiments were run with vegetation dynamics off to maintain the specified canopy composition, structure and soil type combinations, which might not naturally occur.

**Tab. 1:**
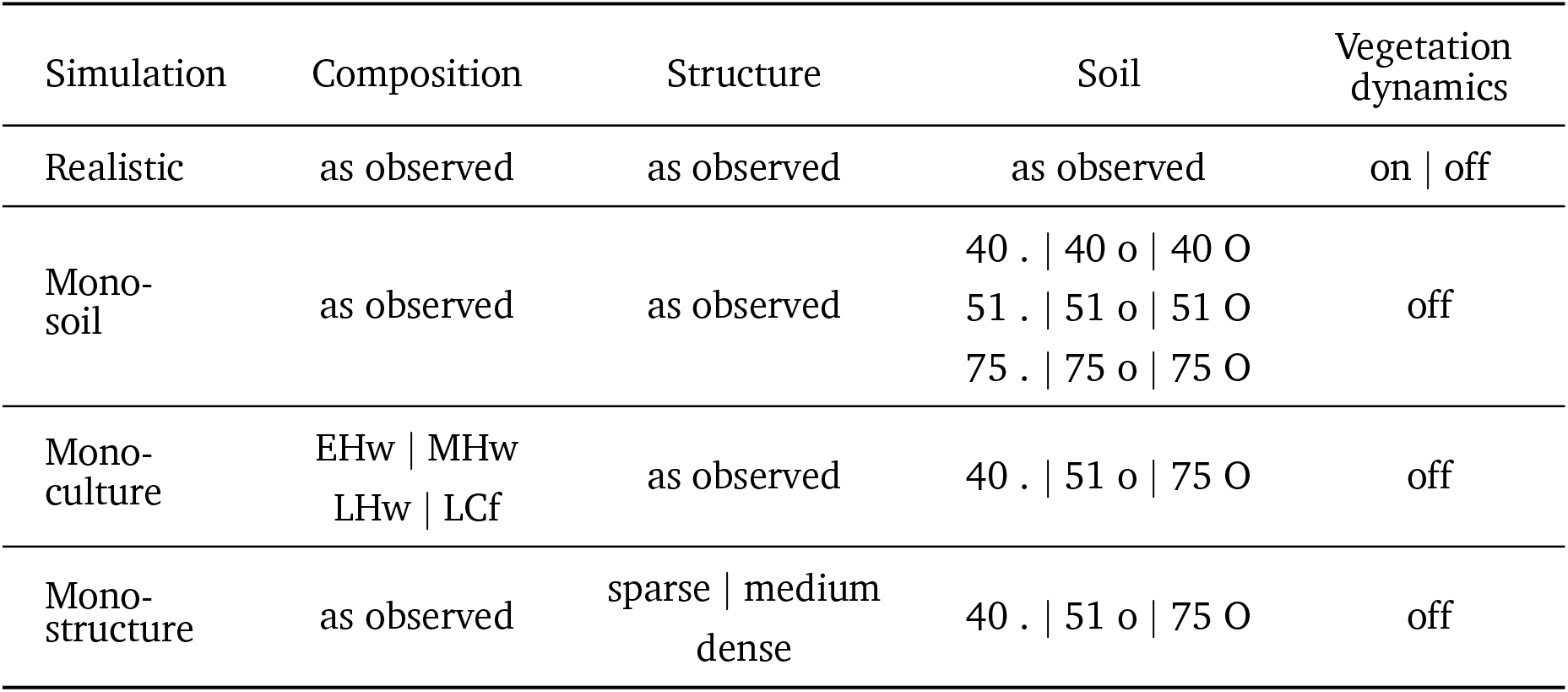
Four sets of simulations run using ED2 on five regions each comprised by three 1-ha sites (15 total) to produce monthly outputs for the years 2006–2015. Multiple entries separated by *|* represent multiple simulations in a set. Mono-soils were simulated at three depths (40, 51, 75 cm) and three textures (.-clayey loam, o-loam, O-sandy loam), and the combinations thereof. Monocultures were simulated for early-, mid- and late-successional hardwoods (EHw, MHw, LHw) and late-successional conifers (LCf). Mono-structures were simulated for sparse, medium and dense canopies.

| Simulation | Composition | Structure | Soil | Vegetation dynamics |
| --- | --- | --- | --- | --- |
| Realistic | as observed | as observed | as observed | on \| off |
| Mono-soil | as observed | as observed | 40 . \| 40 o \| 40 O | off |
|  |  |  | 51 . \| 51 o \| 51 O |  |
|  |  |  | 75 . \| 75 o \| 75 O |  |
| Mono-culture | EHw \| MHw<br>LHw \| LCf | as observed | 40 . \| 51 o \| 75 O | off |
| Mono-structure | as observed | sparse \| medium<br>dense | 40 . \| 51 o \| 75 O | off |

### Forest functional traits, diversity and productivity

Forest functional traits were calculated as patch-averages at 10*×*10-m spatial grain from cohort-level ED2 model outputs of summer 2006. The selected traits represent important ecological and model variables for ecosystem functioning, relating to the distribution of leaf area (LAI), tree diameter and density (basal area, BA), vertical canopy layering (foliage height diversity, FHD), maximum leaf Rubisco carboxylation rate (VC_max_) and specific leaf area (SLA). The physiological traits VC_max_ and SLA were derived from the model values assigned to each PFT and within-patch PFT-mixtures by calculating weighted averages based on LAI and leaf biomass of each cohort. The morphological traits BA and LAI were also derived as patch averages, and FHD was calculated from cohort LAI values following Schneider *et al*. (2017). We then calculated multi-dimensional functional diversity from those traits, specifically morphological and physiological richness, divergence and evenness at 20*×*20, 30*×*30, 40*×*40, 50*×*50 and 100*×*100-m spatial extent following Schneider *et al*. (2017). We defined functional richness as the average of morphological and physiological richness. For forest productivity, we used cohort-level outputs to calculate monthly GPP per 10*×*10-m forest patch and average annual and decadal GPP of 2006–2015.

### Statistical analyses

To study the effects of morphological and physiological diversity on GPP, we fitted a linear model (Matlab 2019a, fitlm) with functional diversity metrics (log(richness), divergence, evenness) as explanatory variables and log(GPP) as the response variable; and a general linear model (Matlab 2019a, fitglme) with soil depth*texture combinations and functional diversity metrics as fixed effects. We also analyzed the model outputs with a random forest model to show the relative feature importance of average functional traits and diversity for predicting GPP at 20*×*20 and 100*×*100-m spatial resolution. Predictor importance was estimated from permutations of out-of-bag predictor observations of 1000 regression trees (Matlab 2019a, fitrensemble, oobPermutedPredictorImportance; Schneider *et al*., 2020), and scaled relative to the fraction of variance explained (*r*^2^) by the model.

We analyzed the effects of morphological and physiological richness on productivity in model experiments (mono-soil, monoculture, mono-structure). We fitted general linear models (Matlab 2019a, fitglme) with log(GPP) as response variable, and soil depth*texture combinations and log(richness) as fixed effects in mono-soil, soil depth*texture, PFT and log(richness) in monoculture, and soil depth*texture, structure and log(richness) in mono-structure experiments, respectively.

## Results

Integrating high-resolution remote observations of forest composition and structure (Fig. S11) into the terrestrial biosphere model ED2 allowed us to predict the field-measured size-structure as basal area per diameter class with an *r*^2^ of 0.94, ranging from 0.70 to 0.93 for individual plant functional types at the fluxtower plot (Fig. S12). This allowed us to study the relationships between morphological and physiological diversity and forest productivity at various spatial scales (20–100-m) along an environmental gradient (5 soil types) in a realistic setting and in model experiments. We first evaluated the model performance by comparing simulated carbon fluxes with eddy-covariance-based estimates at the temperate mixed forest’s fluxtower over the course of ten years, which built the basis for the subsequent diversity–productivity analyses.

### Model evaluation

We found that the model was able to accurately predict the carbon dynamics and seasonality of the Laegern temperate mixed forest averaged across the fluxtower footprint, explaining 86% and 82% of monthly and diurnal variations in GPP as estimated at the fluxtower using eddy-covariance data from 2006 to 2015. Figure 3 shows the simulated carbon dynamics with respect to the most important meteorological drivers and fluxtower estimates. The model captured the seasonality of monthly GPP, but was sensitive to water available to the plants, potentially overestimating their physiological response.

**Fig. 3:**
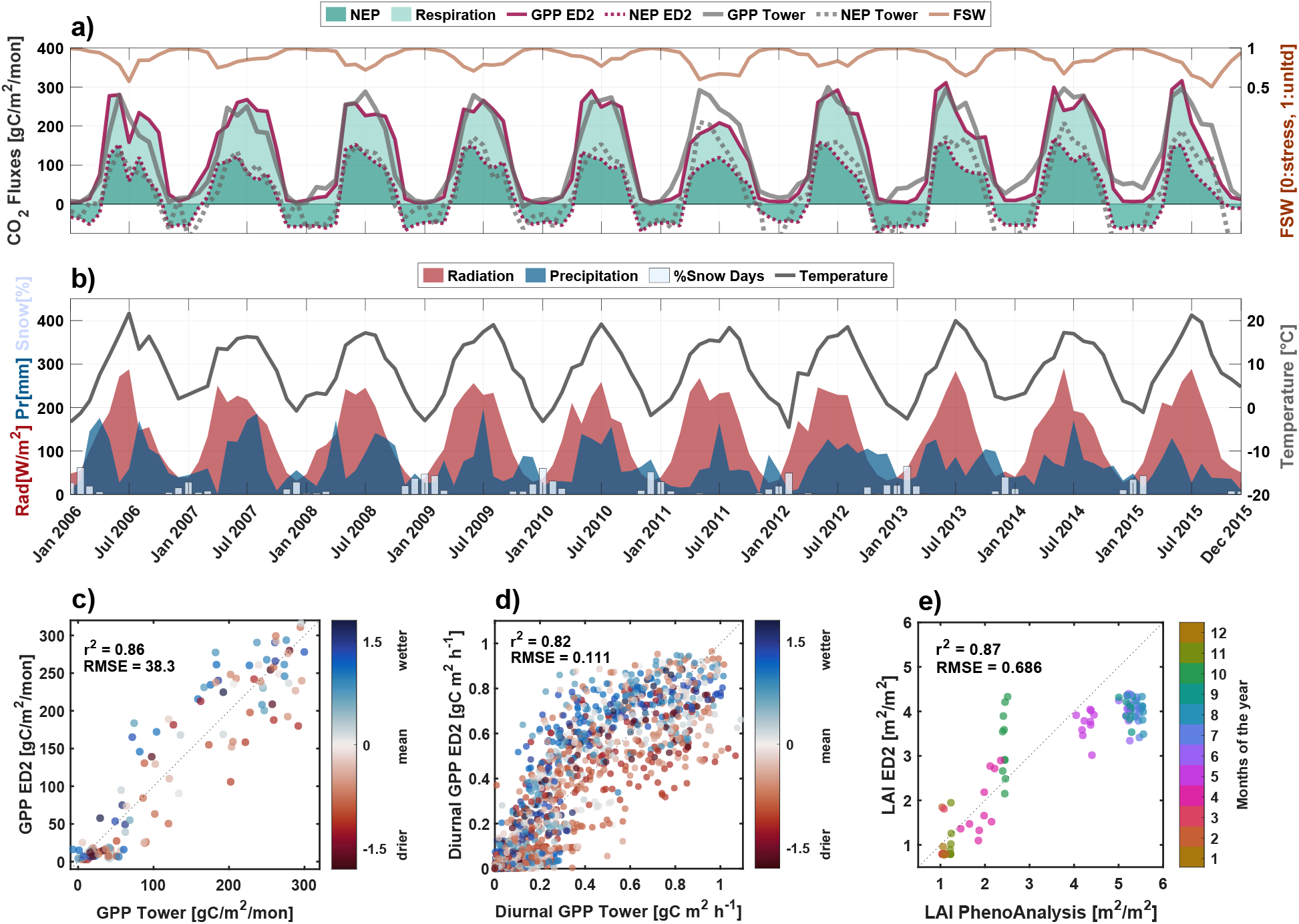
ED2 model evaluation of monthly and diurnal carbon fluxes and monthly leaf area index (LAI) against eddy covariance tower-based and satellite-based estimates in the context of meteorological drivers. The panels show (a) monthly simulated gross primary productivity (GPP, solid red line), heterotrophic and plant respiration (light green area), net ecosystem productivity (NEP, dark green area, dotted red line), the fluxtower estimates of GPP (solid gray line) and NEP (dotted gray line); (b) monthly radiation (red area), precipitation (blue area), percent days with snow cover (white bars), and air temperature (solid gray line); (c) modeled GPP (ED2) against fluxtower estimates (Tower) per month and (d) as monthly average diurnal cycle per hour; (e) modeled LAI (ED2) as monthly average of site-level LAI against satellite-based LAI using PhenoAnalysis to retrieve LAI from Moderate Resolution Imaging Spectroradiometer (MODIS). The colorscale of wetter to drier months is based on a z-score transformation of monthly precipitation values from 2006–2015 to a mean of 0 and a standard deviation of 1 for each month.

For example, annual GPP of the year 2011 was underestimated by 27% (Fig. 3a) following an exceptionally dry spring with precipitation 63% below the decadal average (Fig. 3b). The increased water stress was also reflected by the model parameter FSW (ecosystem-scale down-regulation factor for photosynthesis due to limited soil water availability; FSW = 1 means no soil water stress), which showed values 13% below the growing season average, indicating increased drought stress. Similar effects of reduced GPP and FSW were observed for dry summer months in 2006, 2008 and 2014, and an early GPP drop-off in dry and hot summers of 2013 and 2015. In 2006, the wet spring and fall (115% and 24% above average precipitation) led to 23% higher annual GPP than estimated by the tower despite hot and dry June and July. In summary, the model overestimated tower GPP under wetter conditions and underestimated it under drier conditions, at both monthly (Fig. 3c) and diurnal scales (Fig. 3d), but generally captured the dynamics with little overall bias. Seasonality was generally well predicted, with a slight tendency to an extended productivity at the end of season. This is also reflected in a higher LAI in October in most years due to ED2 predicting delayed leaf fall in comparison with satellite-based LAI (Fig. 3e).

Here, we focus on GPP, but the model shows a good overall performance in simulating carbon fluxes including net ecosystem productivity as estimated by the fluxtower (*r*^2^ = 0.76), even though respiration is difficult to estimate from fluxes at this particular site due to the steep slope, dense upper canopy and potential below canopy CO_2_ drainage (Paul-Limoges *et al*., 2017).

### Diversity–productivity across sites

We found generally positive relationships between functional diversity and productivity across sites, spatial scales, and functional diversity indices when modeling GPP with vegetation dynamics and observed soil types. A general linear model showed highly significant effects of local morphological and physiological richness (p *<* 0.001) and soil types (p *<* 0.001) on average GPP at 20*×*20-m to 50*×*50-m scale, when controlling for different soils between regions (Figs. 4a,b,d,e, S13; Tabs. S2, S3). There is a strong separation in GPP between regions 1-2 and the rest, following differences in soil matric potential and depth (Fig. 4i). Functional divergence and evenness effects were less strong and more variable from positive to neutral (Figs. S14, S15, Tabs. S4, S5, S6, S7).

**Fig. 4:**
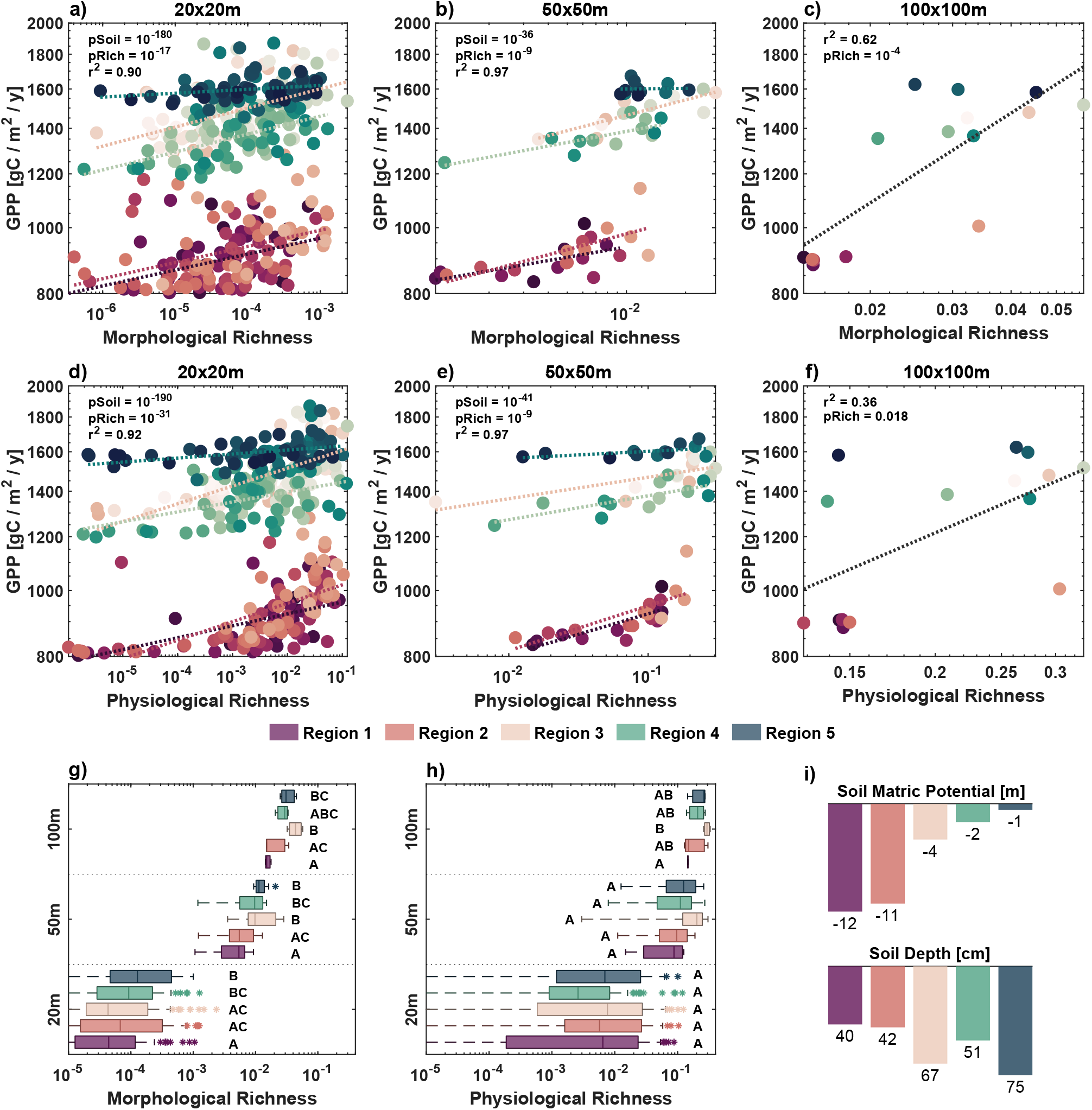
Relationship between (a-c) morphological and (d-f) physiological richness and productivity as average annual gross primary productivity (GPP), simulated across five regions over the course of a decade with vegetation dynamics at (a,d) 20×20-m, (b,e) 50-50-m, and (c,f) 100×100-m scale. Statistics in (a,b,d,e) are based on a general linear model controlling for different soils between regions, while (c,f) shows a linear regression across all sites. Panels (g,h) show the overlap of richness for each region across spatial scales. Significant differences between the regions within three spatial scales are indicated by the compact letter display based on Fisher’s least significant difference test. (i) shows soil matric potential as indicator of soil water available to the plants (more negative numbers mean more strength required to extract water), and corresponding soil depth of the five regions. Colors are consistent across panels and refer to the five different regions and their unique soil properties.

At 100*×*100-m scale, we found significant positive relationships with productivity across sites for morphological (p *<* 0.001) and physiological (p *<* 0.05) richness (Fig. 4c,f). Figure 4g,h shows the increase of functional richness and narrowing of the range with spatial extent. Morphological richness was significantly different between regions with the shallowest and deepest soils (Fig. 4g), while physiological richness did not differ significantly except between region 1 and 3 at 100*×*100-m (Fig. 4h).

A random forest model analysis showed that environmental variables related to soil water were the dominant predictors of GPP at 100*×*100-m scale, but functional diversity still showed a relative importance of 9–18% and up to 17–20% for predicting GPP when considering indirect effects through correlation to other functional traits (Fig. 5). At local scale, community-weighted average physiological traits had the highest relative importance (51%) when predicting GPP with vegetation dynamics enabled, whereas community-weighted average morphological traits were most important (27%) when vegetation dynamics was disabled (Fig. 5). In both cases, the importance of functional diversity strongly increased when both direct and indirect effects were considered (11–55% with vegetation dynamics on, 9–26% with vegetation dynamics off). Functional richness was the overall most important diversity metric, and both morphological and physiological richness were important and varied significantly with GPP (Figs. 4, 5).

**Fig. 5:**
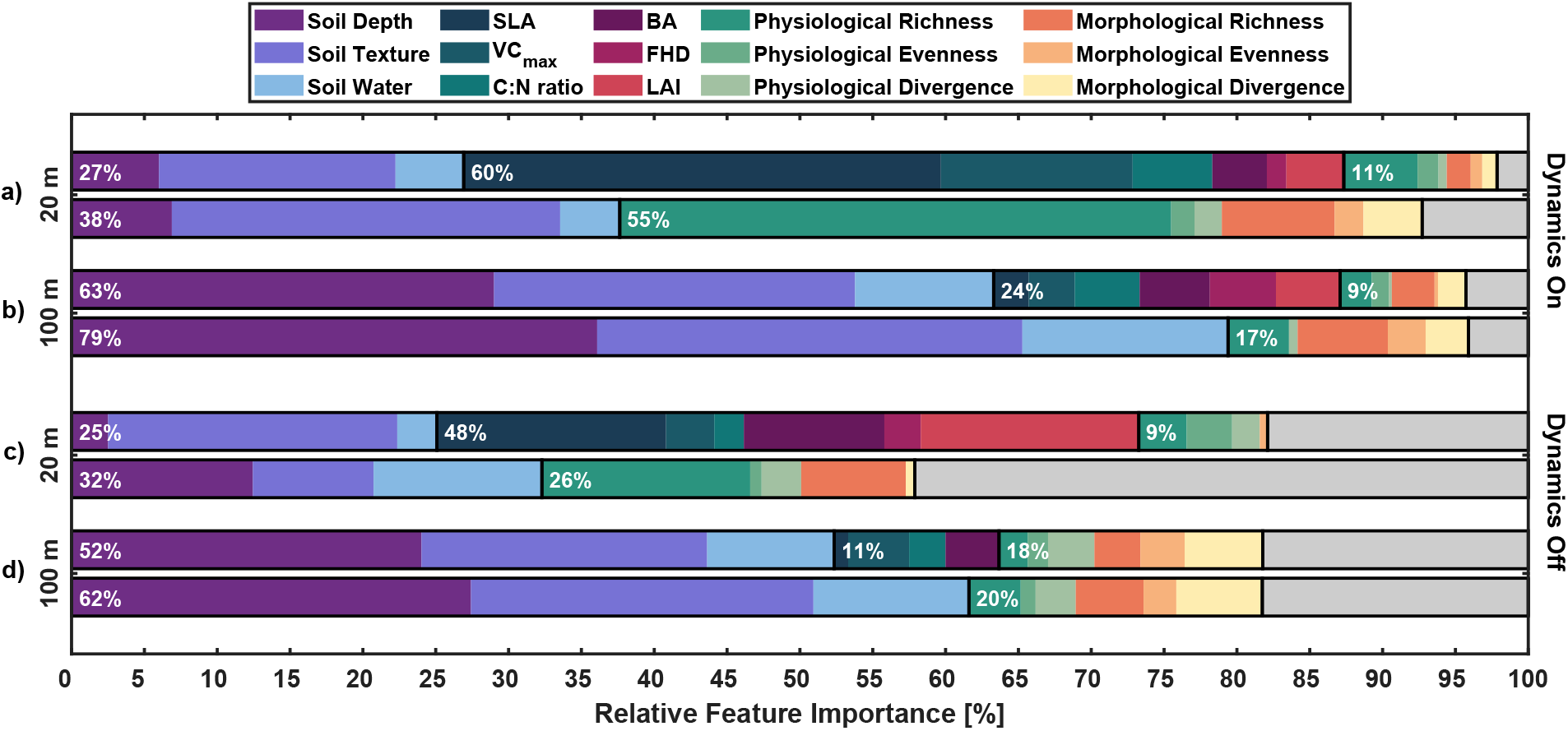
Random forest feature importance to predict average annual gross primary productivity, simulated with vegetation dynamics on (a,b) and off (c,d) at 20 m (a,c) and 100 m (b,d) spatial scale, scaled by the total variation explained (*r*^2^) by the model. The gray bars indicate unexplained variance. The white numbers indicate the relative importance of a group of variables, while the partitioning within the group has to be interpreted with care due to possibly stronger correlations among the different explanatory variables in a group than between the groups. By including (top of each spatial scale) and excluding (bottom of each spatial scale) average functional traits, we can see direct and indirect effects of functional diversity on productivity.

### Diversity–productivity model experiments

In natural ecosystems, co-varying drivers of diversity and productivity, such as different soil characteristics, make it difficult to draw conclusions about diversity effects on productivity (Loreau, 2000; Grace *et al*., 2016). The model environment enabled us to run experiments with homogeneous soils (mono-soils), monocultures and mono-structures to disentangle the effects of soil, morphological and physiological richness on long-term productivity.

We found that functional richness, as average of morphological and physiological richness, on mono-soils had a significant positive effect on GPP (p *<* 0.001) at local scale (20–30 m, see Fig. 6a, Tab. S8). Only when considering productivity per leaf area, the richness effect remained strongly positive among all scales (p *<* 0.001 at 20–100 m, Fig. S16, Tab. S9). Evenness and divergence showed mixed results from positive to negative and highly to non-significant relationships without clear patterns (Fig. S17).

**Fig. 6:**
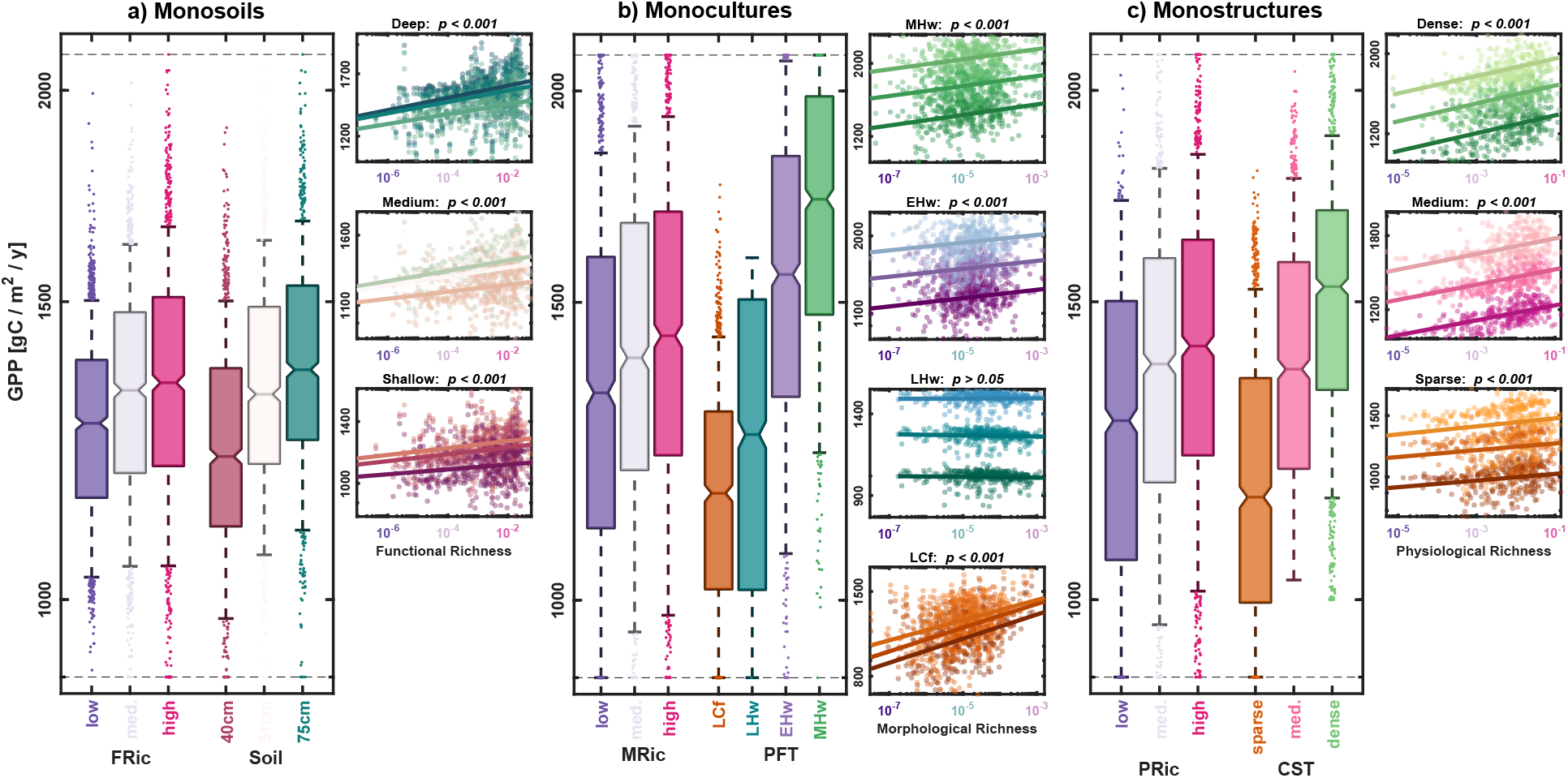
Boxplots comparing low, medium and high (a) functional (FRic), (b) morphological (MRic) and (c) physiological richness (PRic) with (a) shallow, medium and deep soil, (b) late-successional conifers (LCf), late-, early- and mid-successional hardwoods (LHw, EHw, MHw) plant functional types (PFT), and (c) sparse, medium and dense canopy structural types (CST), respectively, at 20*×*20-m spatial extent. Simulations were run with single soil type (mono-soil, each 375 areas at 20*×*20 m), single PFTs (monoculture, 375 *×* 3 soils), and CSTs (mono-structure, 375 *×* 3 soils). The scatter plots show all data points and linear regression lines for (a) three soil textures (clayey loam, loam, sandy loam) per soil depth, (b) three soils per PFT and (c) three soils per CST. The three soils were clayey loam at 40 cm, loam at 51 cm, and sandy loam at 75 cm depth. P-values are shown based on general linear models controlling for soil.

The monoculture simulations further showed that the effect of morphological richness on productivity was highly significant at local scales (p *<* 0.001 at 20–40 m, Tab. S10), but the effect varied by PFT, with no effect for late-successional hardwoods and the most positive effect for late-successional conifers (Figs. 6b, S18). A positive effect was also found for physiological richness across canopy structures (Fig. 6c; Tab. S11), with strongest positive effect for tall and dense canopies at scales from 20–50 m (Fig. S19).

In all three experiments, the maximum difference in productivity between the lowest and highest productivity soils, plant functional types and canopy structures, respectively, was larger than between lowest and highest forest functional diversity (Fig. 6). Also, there was no diversity effect at 100*×*-m scale in the experiments (Figs. S16, S18, S19), indicating that morphological and physiological diversity were especially important at local scale.

## Discussion

Our results show that there are significant positive effects of morphological and physiological diversity on average annual forest productivity from 2006–2015 at local spatial scales (tens of meters), and that this phenomenon holds both within soil types and across an environmental gradient. These positive diversity effects are likely stronger in reality, because terrestrial biosphere models do not include all processes and interactions between species and forest patches, such as the plastic responses of species traits to environmental conditions and stand diversity (Roscher *et al*., 2018). At larger spatial scales (100*×*100-m), a positive correlation between functional diversity and productivity occurs along the environmental gradient of differing soil textures and depth, comparable to Durán *et al*. (2019). This effect is more complex, however, because soils may act as confounding factors as evidenced by the loss of the hectare-scale diversity–productivity relationship when simulating uniform soil types. In the following sections, we discuss the processes and mechanisms in the terrestrial biosphere model that drive the influence of morphological, physiological and environmental aspects of the forest ecosystem on its functioning.

### Morphological diversity and productivity

Morphological diversity of the plant canopy structure controls the radiative transfer and light distribution throughout the canopy, with higher diversity in canopy structure leading to a more even distribution of light throughout the canopy and higher light use efficiency and productivity (Ishii *et al*., 2004; Williams *et al*., 2017; Kükenbrink *et al*., 2021). In the ED2 model, higher morphological diversity also relates to higher complementarity in soil water use, as plant size (i.e. tree diameter and height) determines the plant rooting depth. High local variability in plant size means high variability in rooting depth, thus reducing competition for water in the same soil layer, increasing complementarity and overall productivity through reduced stress (Loreau *et al*., 2001).

The effect of morphological diversity on productivity is likely underestimated in the model, since interactions between forest patches, such as changed radiative transfer through shading or sun exposure from neighboring patches, were not modeled. However, the remotely sensed structural information that we used as initial conditions in the model likely did integrate processes that are not modeled explicitly, such as nutrient cycles, wind throw disturbance or forest management. Morphological diversity effects in the model are therefore mostly stemming from observed relationships between diversity and other stand-level structural properties that influence model productivity, such as increased stem density and basal area in structurally more diverse forest areas as observed in other studies (Barrufol *et al*., 2013; Morton *et al*., 2016). Such a limitation is common in terrestrial biosphere models (Fisher & Koven, 2020) and shows the importance of integrating empirical observations of forest structural variability into the models (e.g., Caylor *et al*., 2004; Hurtt *et al*., 2010; Fischer *et al*., 2016; Levine *et al*., 2016; Longo *et al*., 2018; Braghiere *et al*., 2019). Additional efforts are needed to map forest structure and composition repeatedly and to operationalize data-model integration and model benchmarking over time.

Finally, while tree diameters and density can vary freely and be integrated into the model as initial conditions, other structural variables such as height or biomass are defined by static PFT-specific allometric relationships. ALS measurements show, however, a high degree of variability in diameter-height relationships. Incorporating plastic allometry is an interesting area for future model development, since tree heights (and many other variables of canopy structure) can be derived from laser scanning at different spatial scales (e.g. Schneider *et al*., 2014, 2019, 2020).

### Physiological diversity and productivity

The diversity of physiological traits (here only between PFTs, within PFTs these traits are assumed to be constant) represents the variety of functional strategies and life forms (Reich *et al*., 2003; Díaz *et al*., 2015), influencing an ecosystem’s ability for adaptation, defense and recovery that will determine its long-term stability and productivity (Díaz & Cabido, 2001; Mason & de Bello, 2013). There are two main effects causing productivity to increase as physiological diversity increases that are captured in the model: (1) a temporal stability and insurance effect, and (2) a selection and dominance effect driven by the physiological properties of the model PFTs.

With respect to (1), the temporal asynchrony of plant communities, species or functional types has been shown to positively influence ecosystem productivity and stability (Tilman *et al*., 2014; Loreau & de Mazancourt, 2013; Craven *et al*., 2018), and this effect is also represented by ED2. For example, late-successional conifers can photosynthesize earlier and later in the growing season during leaf development and senescence of deciduous species, and their productivity is more stable due to lower stomatal conductance that leads to lower drought sensitivity (Fig. 7a). In contrast, deciduous hardwoods are more productive during peak greenness, but react more strongly to meteorological variability (Fig. 7a). The ED2 model’s plant functional types represent the different seasonal dynamics and stress responses that can explain positive diversity-effects at decadal temporal scales, consistent with previous ecological modeling experiments (Cardinale *et al*., 2004).

**Fig. 7:**
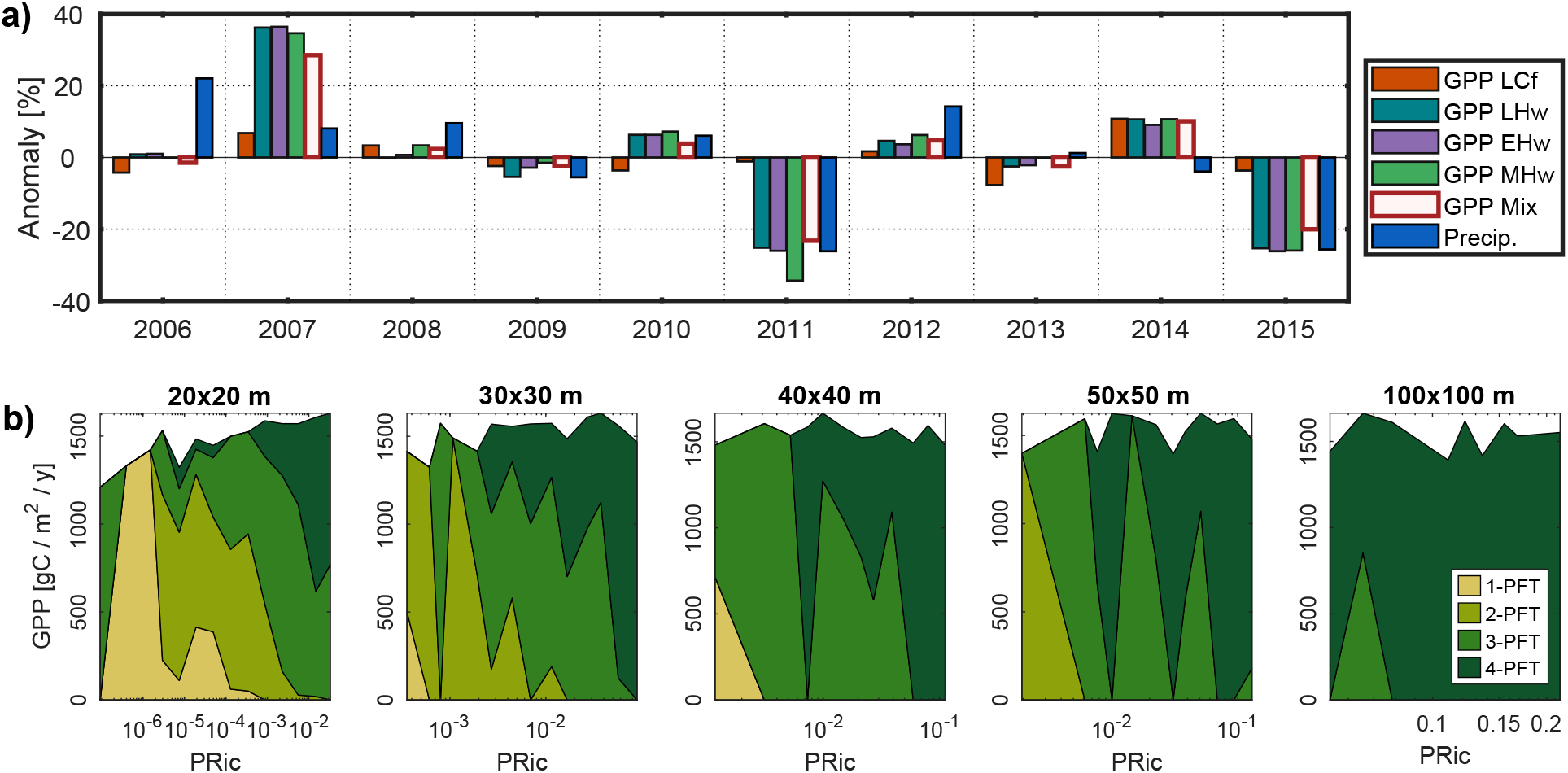
Annual variability in gross primary productivity (GPP) of plant functional type (PFT) monocultures and mixtures, and contributions of PFT-mixing to physiological richness (PRic) and GPP at different spatial scales. a) Annual relative anomaly compared to the decadal mean GPP of late-successional conifers (LCf), late-, early- and mid-successional hardwoods (LHw, EHw, MHw), simulated in monocultures, compared to simulations using observed PFT-mixtures (Mix) on a 51-cm loamy soil. The blue bars shows annual precipitation anomalies for context. b) Cumulative contributions of mixtures of 1-4 PFTs to GPP by physiological richness across spatial scales from 20*×*20-m to 100*×*100-m extent. PRic shows saturation towards larger scales, where almost all PFTs are present at all sites, even though in different abundances.

With regard to (2), there can be a selection and dominance effect in which dominance by species or plant functional types with particular traits affects ecosystem functioning (selection effect; Loreau *et al*., 2001). As in most central European temperate forests, in our study area beech is the dominant late-successional hardwood species. We modeled late-successional hardwoods with lower VC*_max_* and higher SLA to represent their lower maximum productivity and higher shade tolerance compared to early- or mid-successional hardwoods (Peters, 1992; Heiri *et al*., 2009, see also Fig. 6). The consequence is that early- and mid-successional hardwoods have higher productivity if enough light is available but cannot maintain high leaf area and productivity in dense canopies or understory, while late-successional hardwoods can maintain dense, tall canopies and productivity under lower light conditions, but with lower maximum productivity. Therefore, lower-diversity beech forests will have lower productivity than mixed forests with local disturbance and location-specific growing conditions that allow early and mid-successional species to compete, co-exist and be highly productive. This diversity of plant community and structure was captured by the integration of remote observations into the model that capture different development stages and the effect of local disturbance processes, which is characteristic of productive and sustainable forests over time (Cardinale *et al*., 2004; Silva Pedro *et al*., 2016; Dolezal *et al*., 2020).

Overall, physiological diversity effects may also be underestimated in the model, since physiological complementarity in nutrient use, complementairty in soil microbial communities and related co-benefits of microbial diversity, or complementarity in insect defence strategies were not included in the model formulation. In addition, physiological traits were defined by the plant functional type of each tree species, which limits functional diversity in physiological traits to mixtures of typically three to four PFTs, even when the total species richness may be considerably higher (13 tree species at our study site). Opportunities exist to implement more flexible traits, for example in terms of within-canopy trait plasticity with light availability (Berzaghi *et al*., 2020; Xu & Trugman, 2021). Further development is needed, for example towards flexible traits such as leaf mass per area, nitrogen or V*_cmax_*, as proposed by Sakschewski *et al*. (2015) and adapted to European forests by Thonicke *et al*. (2020). The combined ability to represent plastic trait variation and integrate it from remote remote sensing could greatly advance the representation of forest functional diversity (e.g. Schneider *et al*., 2017; Durán *et al*., 2019; Wang *et al*., 2020) and the modeling of carbon fluxes and responses to climate change (Schimel *et al*., 2019).

### Spatial scale and environmental drivers

Our results showed that the diversity effects discussed above primarily act at the local (sub-hectare) spatial scale, where functional diversity varies within a large range from low to medium diversity (Fig. 4g,h). This is related to the community structure of the forest, which has high alpha- and low beta-diversity (Schneider *et al*., 2017), a situation that is typical for most temperate forests (Swenson *et al*., 2012). Additionally, most plant interactions in forests happen at the scale of tens of meters (Wang *et al*., 2015), and thus our results are consistent with findings from field-based studies (Chisholm *et al*., 2013; Poorter *et al*., 2015; Liu *et al*., 2018). At larger spatial scales, diversity can be saturated when beta-diversity is low. This can lead to functional redundancy (Rosenfeld, 2002) and a weakening of the diversity–productivity relationship (Jochum *et al*., 2020). In our study, a saturation of physiological diversity with increasing spatial extent might have occurred earlier than in reality due to the limited number of plant functional types (Fig. 7b) and negligence of intraspecific physiological diversity present at this site (Guilĺen-Escribà *et al*., 2021).

The positive correlation between diversity and productivity at the hectare-scale was considerably weakened when the sites were simulated with uniform environmental conditions, indicating co-variation of soil properties with plant diversity and productivity. This suggests that the environmental gradient, characterized by variations in soil depth and texture in the model, was the main driver of productivity at this larger scale, limiting plant productivity through limiting resources such as water availability, while functional diversity was saturated. The effect of soil texture and depth on forest productivity may be overestimated because the model shows a higher-than-observed sensitivity of productivity to plant water availability, which is indicated by monthly and diurnal comparisons to the fluxtower (Fig. 3) and annual GPP anomalies following precipitation (Fig. 7a). The steep, fractured limestone bedrock might offer the trees opportunities to access water below the average soil depth, and this currently cannot be represented by ED2. Therefore, the actual trees at Laegern may exhibit more resistance to water stress than our analysis implies.

### Conclusions and outlook

Functional diversity had a significant positive effect on long-term forest productivity at local spatial scales (tens of meters) in terrestrial biosphere simulations in a temperate mixed forest. This result was found across all soil types and including morphological and physiological diversity, with strongest effects for functional richness. This implies that local functional richness, which can be measured with remote sensing across large spatial extents and in remote ecosystems, is an important driver of productivity in temperate mixed forests. This finding is in line with studies that analyzed species richness-productivity relationships in field plots (Chisholm *et al*., 2013; Poorter *et al*., 2015).

At larger spatial scales (hectare-scale and above), our model experiments showed that diversity’s influence on productivity saturated and that soil depth and texture appeared to be the main drivers of spatial variation in forest productivity at this scale. We note that the influence of soil properties may be overestimated due to the challenge of modeling soil water availability to the plants in this complex terrain. The diversity saturation may in part be due to the community structure of this forest and the limited representation of functional diversity in the model, i.e. physiological diversity being limited to mixtures of plant functional types. We also note that the saturation of biodiversity–ecosystem functioning relationship at the larger scale does not imply that this relationship is not important in driving global forest productivity. That is, if diversity is low at the smaller scale, this still sums up to low productivity at the large scale.

The integration of spatially explicit remotely-sensed trait information in Earth system models is crucial to study the role of biodiversity in carbon cycling and to predict impacts and feedbacks of climate change. The advancement of spaceborne remote sensing will help to characterize plant functional traits and diversity globally, for example morphological diversity using lidar (Schneider *et al*., 2020) and radar (Bae *et al*., 2019) or physiological diversity using imaging spectroscopy (Cawse-Nicholson *et al*., 2021), and will provide new measurements of ecosystem functioning (Schimel *et al*., 2019) to initialize and benchmark global terrestrial biosphere models that represent structurally and functionally diverse ecosystems (Fisher *et al*., 2018).

## Acknowledgements

The research carried out at the Jet Propulsion Laboratory, California Institute of Technology, was under a contract with the National Aeronautics and Space Administration (80NM0018D0004). Government sponsorship is acknowledged. This study was supported by the University of Zurich Research Priority Program on ‘Global Change and Biodiversity’ (URPP GCB). Support was also provided by the NASA HyspIRI Preparatory Activity NNH11ZDA001N-HYSPIRI grant “Linking Terrestrial Biosphere Models with Remote Sensing Measurements of Ecosystem Composition, Structure, and Function” to PRM. ML was supported by the NASA Postdoctoral Program, administered by Universities Space Research Association under contract with NASA. Additional funding was provided by a Swiss Government Excellence Scholarship awarded to VS through the U.S. Fulbright Student Program. Meteorological data was provided by the Federal Office for the Environment (FOEN), the Swiss Federal Laboratories for Materials Science and Technology (EMPA) and MeteoSchweiz (Climap). Stand polygon data was provided by Aargauisches Geografisches Informationssystem (AGIS), Departement Bau, Verkehr und Umwelt, Abteilung Wald (last updated on 27 February 2015) and by Geographisches Informationssystem (GIS-ZH), Amt für Landschaft und Natur, Abteilung Wald (last updated on 16 September 2015). Soil data corresponds to Bodenkarte Baden (Landeskarte der Schweiz 1:25’000, Blatt 1070), provided by Eidgenössische Forschungsanstalt für Agrarökologie und Landbau (FAL).

## Author contribution

FDS, MES, PRM, FM and BS designed research; FDS, PRM, ML, EPL, and VS performed research; FDS, PRM, EPL, and VS analyzed data; and all authors contributed to data interpretation and manuscript writing.

## Data availability

The data that support the findings of this study will be made openly available in a repository (to be done at acceptance of the article in a peer-reviewed scientific journal). The most up-to-date ED2 source code is available at https://github.com/EDmodel/ED2, and the version used for this study is permanently deposited on https://dx.doi.org/10.5281/zenodo.3365659.

The authors declare no conflict of interest.

## Supporting Information

**Fig. S1:**
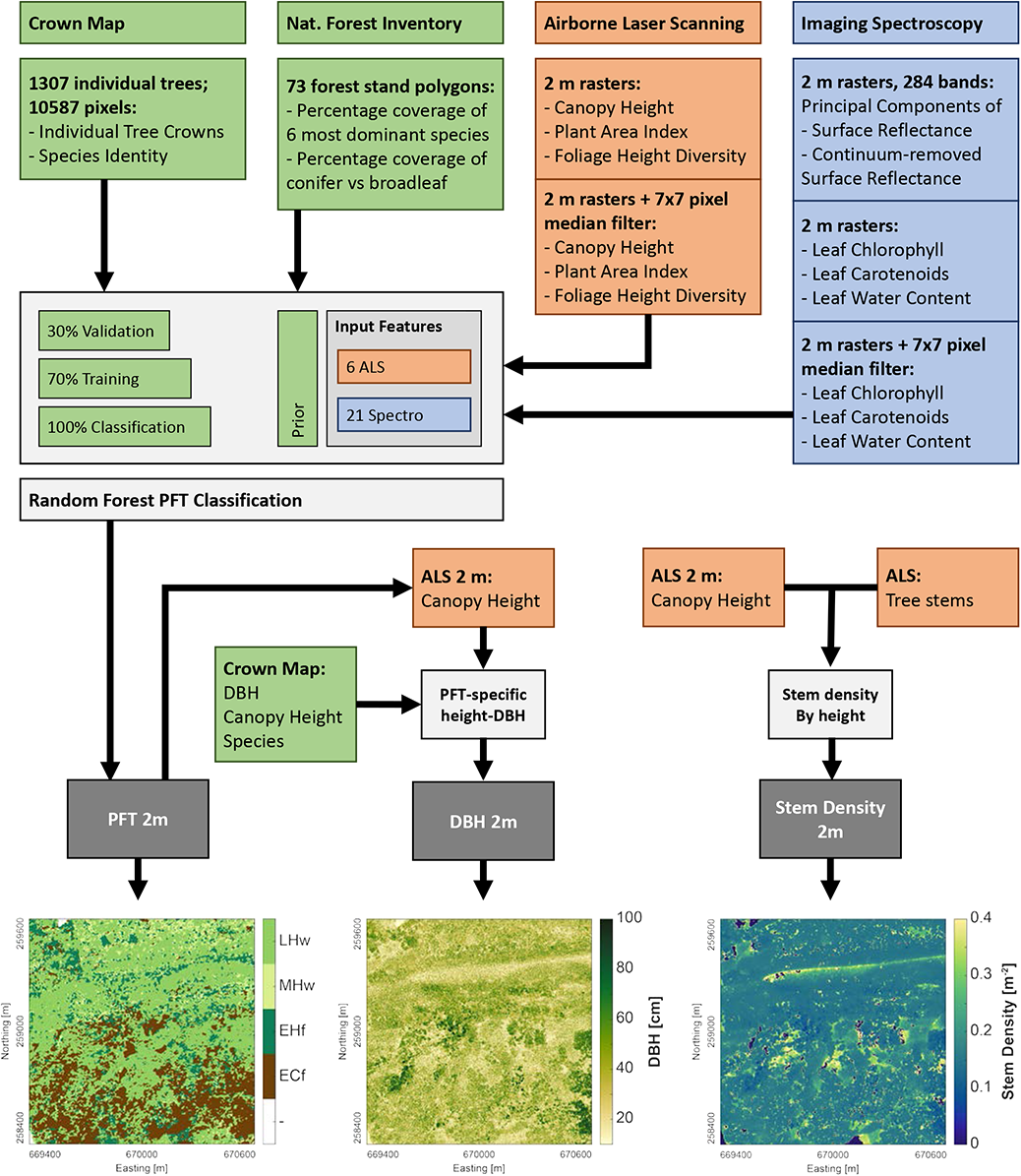
Flow diagram showing the methodology and the remote sensing and field data used to derive high-resolution (2×2-m) maps of plant functional types (PFT), tree diameter at breast height (DBH), and tree stem density. PFT, DBH and stem density were the main variables integrated into the ED2 model to initialize forest composition and structure.

**Fig. S2:**
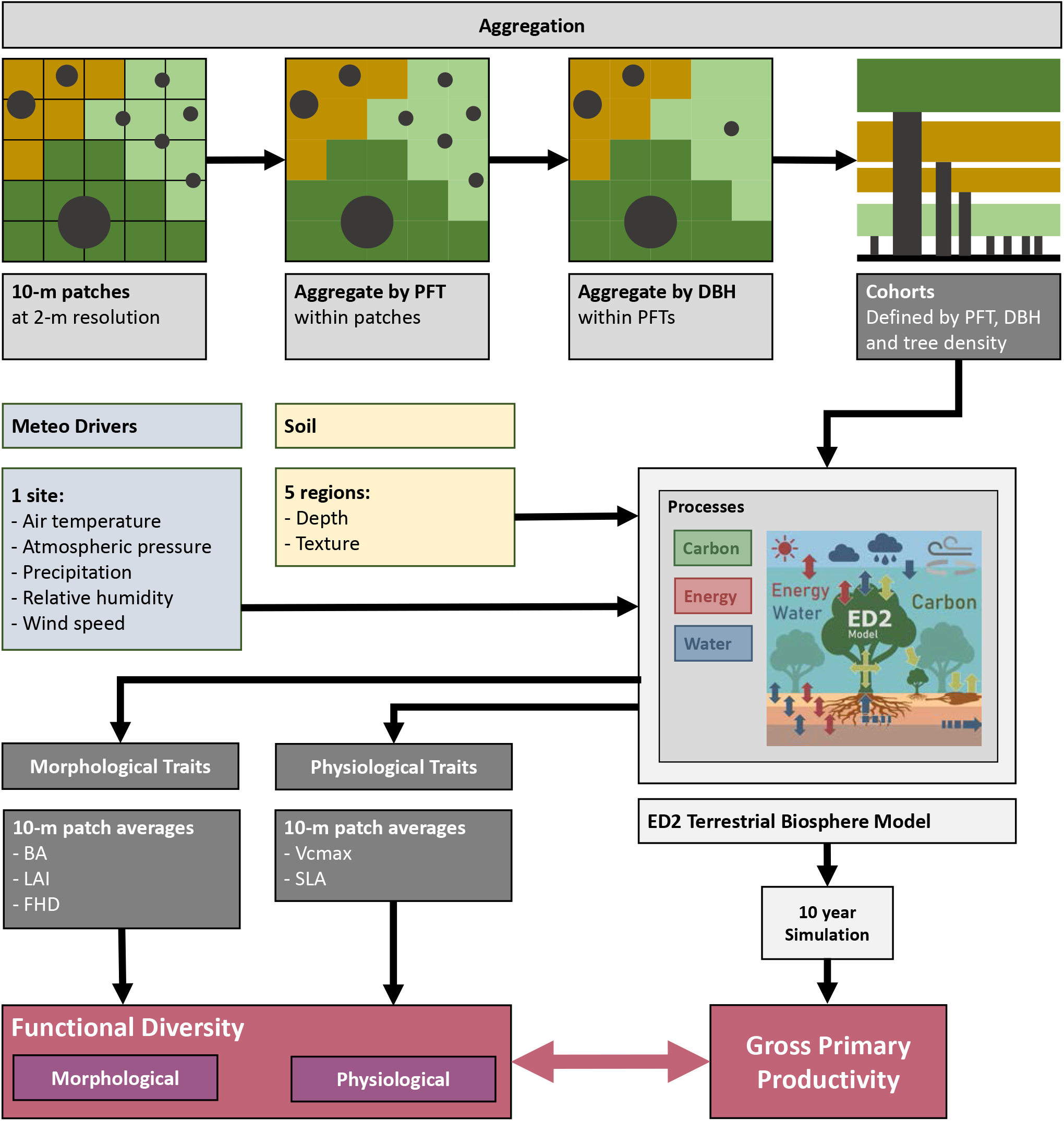
Aggregation of 2-m remote observations of forest composition and structure for integration into the ED2 terrestrial biosphere model as model cohorts within 10×10-m patches. The flow diagram shows the model inputs and outputs that were used to study functional diversity–productivity relationships.

**Fig. S3:**
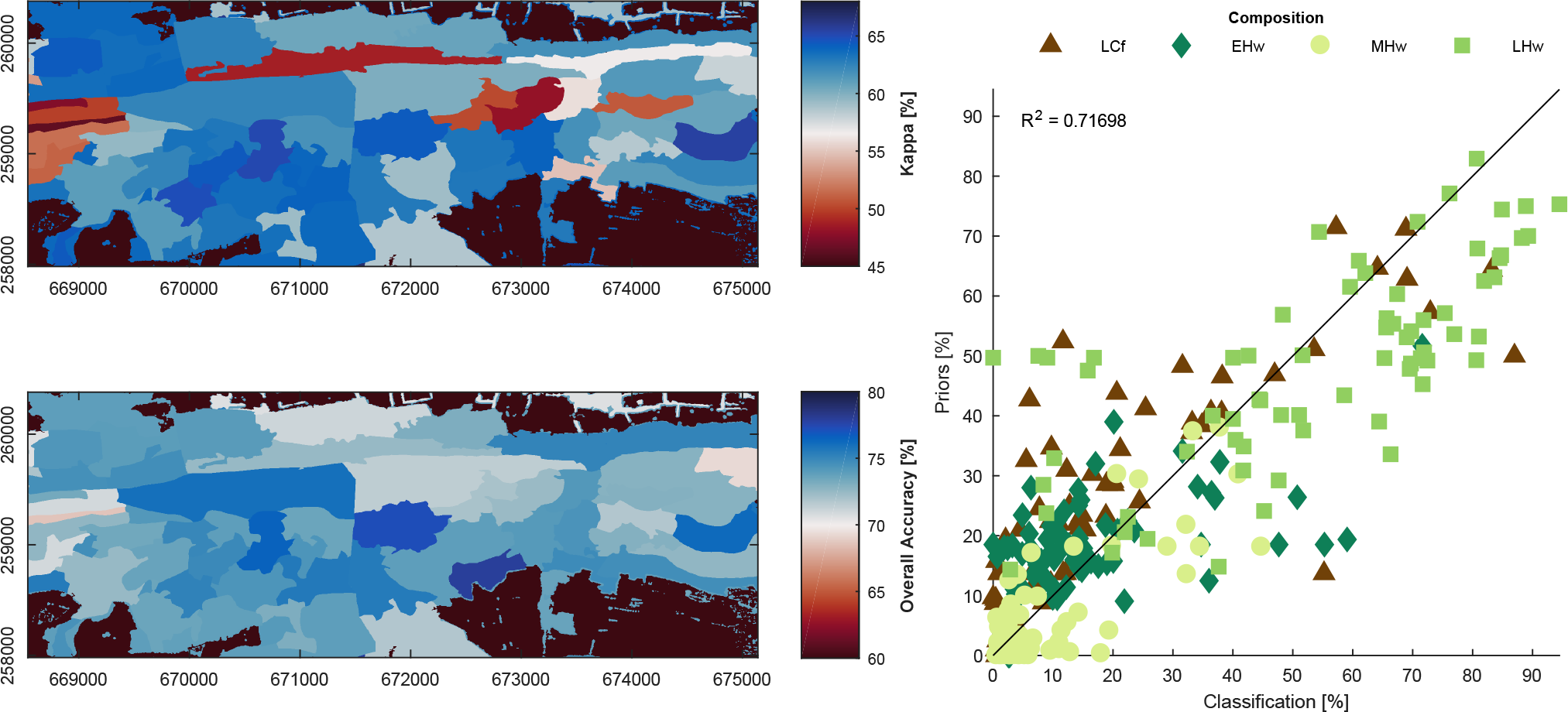
Results of random forest classification of the plant functional types late-successional conifers (LCf), early hardwoods (EHw), mid hardwoods (MHw) and late hardwoods (LHw). The maps show kappa and overall accuracy for each polygon that was separately trained, tested and predicted. The right panel shows how the prior information of composition of plant functional types compares to the actual classification for each polygon.

**Fig. S4:**
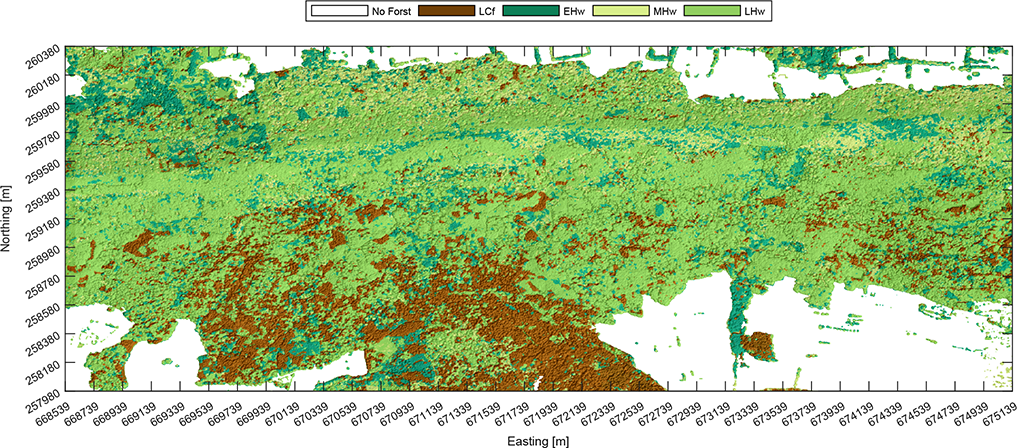
Map of plant functional types classified using ground-based and airborne remote sensing data. The plant functional types are late-successional conifers (LCf), early hardwoods (EHw), mid hardwoods (MHw) and late hardwoods (LHw).

**Fig. S5:**
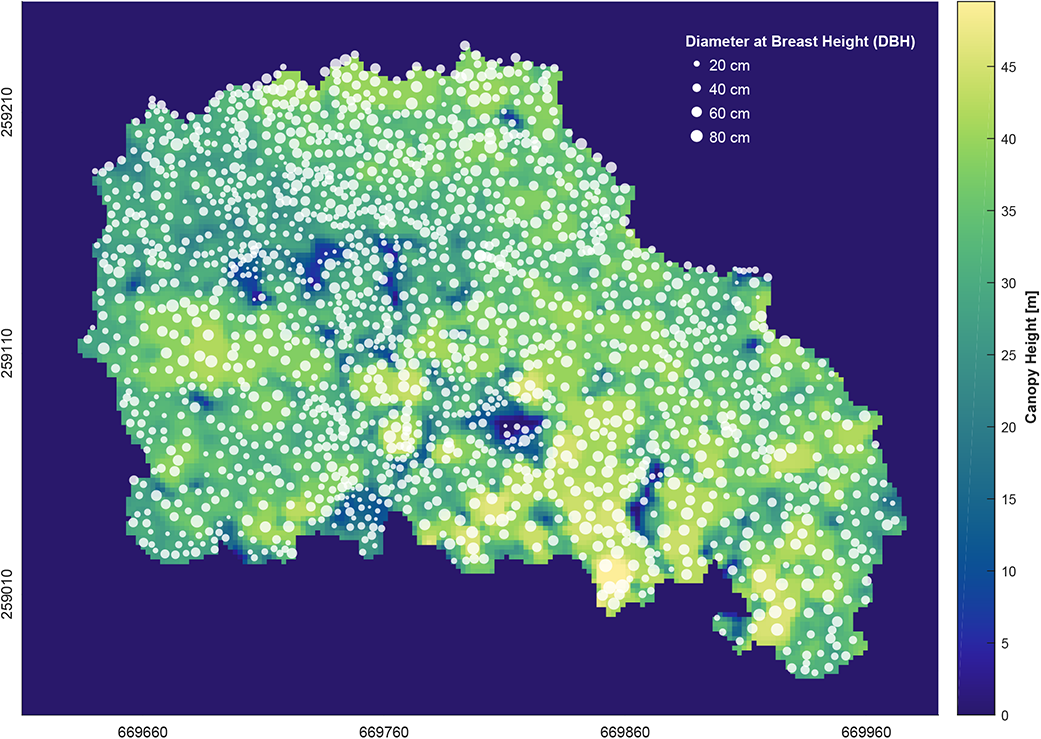
Canopy height and stem distribution as derived from airborne laser scanning data on the 5.5-ha field plot. Canopy height is a continuous raster at 2 m spatial resolution. The white dots represent segmented tree stems with variable size according to the assigned diameter at breast height (DBH), estimated using allometries based on canopy height. The location and distribution of DBH values is not spatially explicit, but should rather represent the correct size distributions of the plot.

**Fig. S6:**
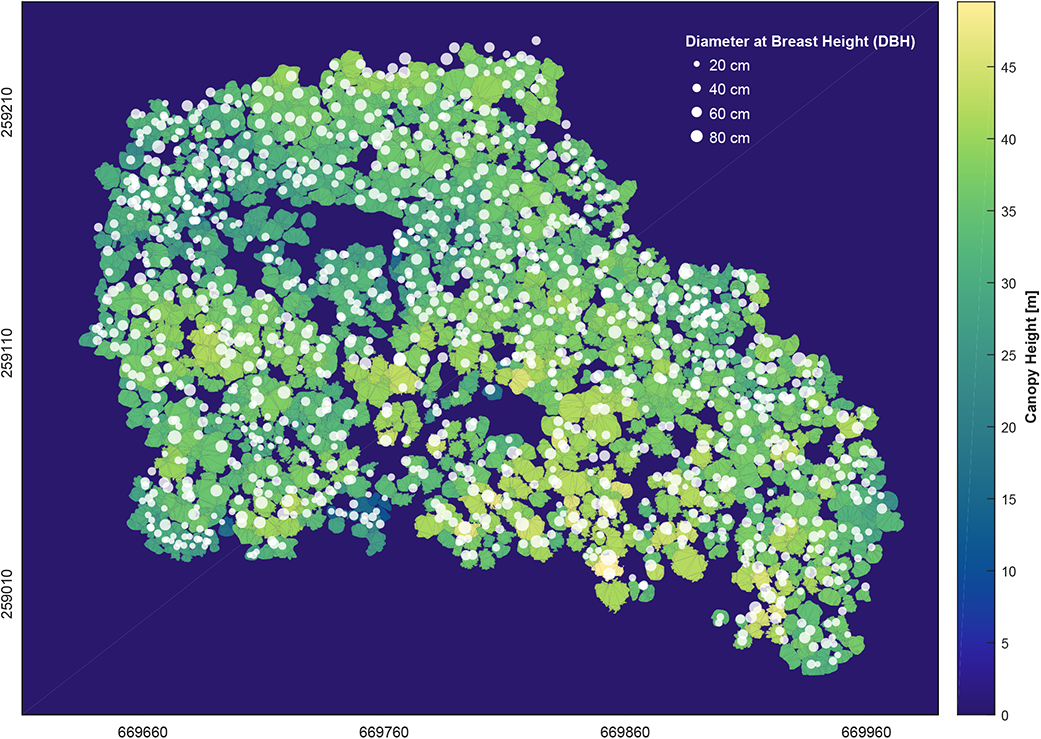
1307 crown polygons colored by canopy height for all dominant and co-dominant trees with a diameter at breast height (DBH) above 20 cm as measured on the 5.5-ha field plot. White dots represent the corresponding tree stem positions with variable size according to the DBH measured in the field.

**Fig. S7:**
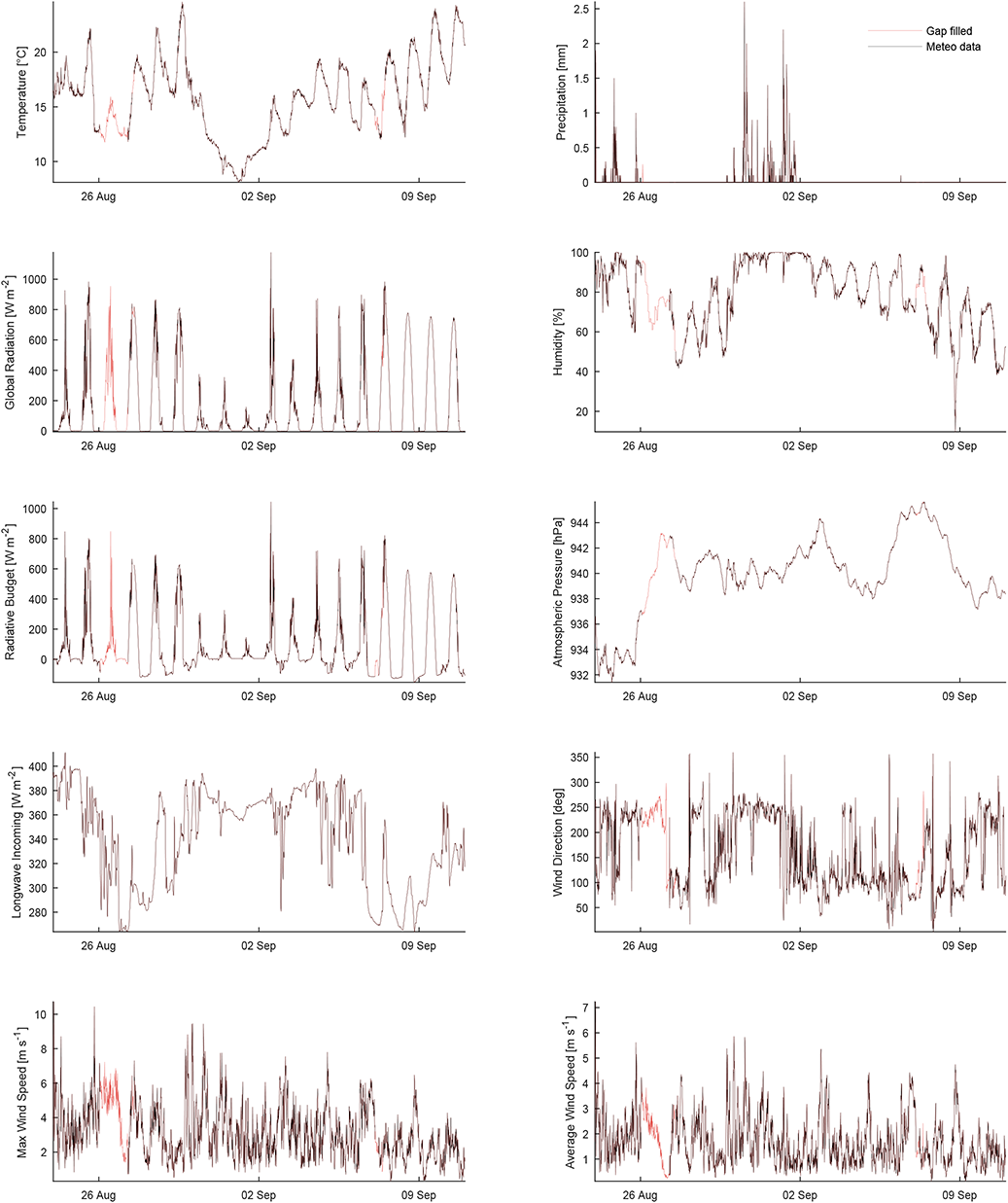
Example of gap-filled meteorological drivers at 10 minutes temporal resolution in the period of 24 August to 10 September 2012. The gap-filled parts of the time series are displayed in red.

**Fig. S8:**
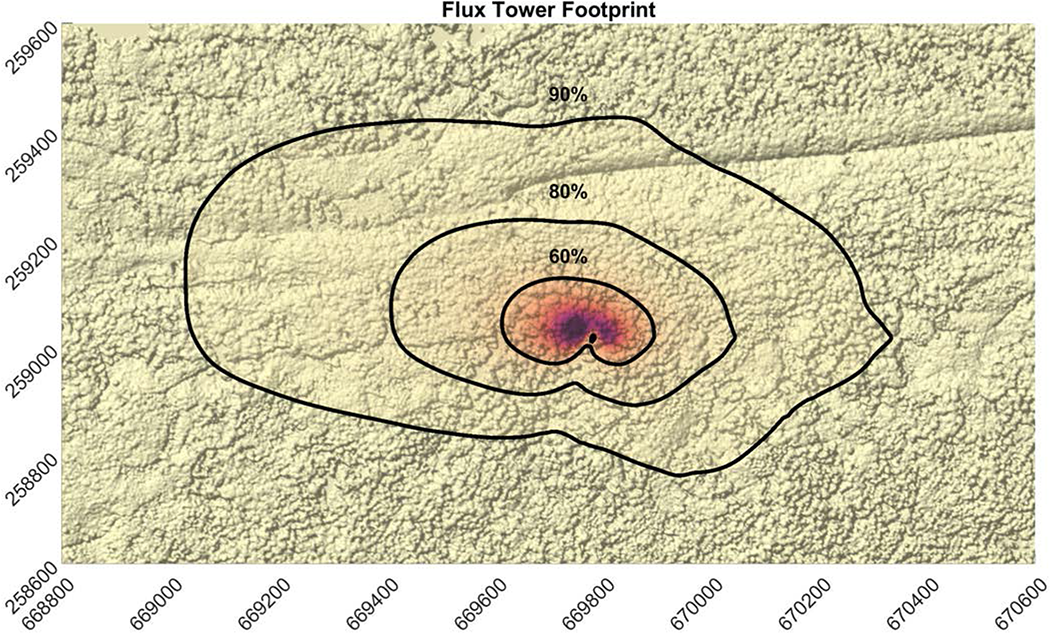
Average daytime fluxtower footprint for the year 2016, with darker colors indicating higher flux density. Contour lines are shown for 60, 80 and 90% of the total footprint. The background map shows the shaded canopy height model.

**Fig. S9:**
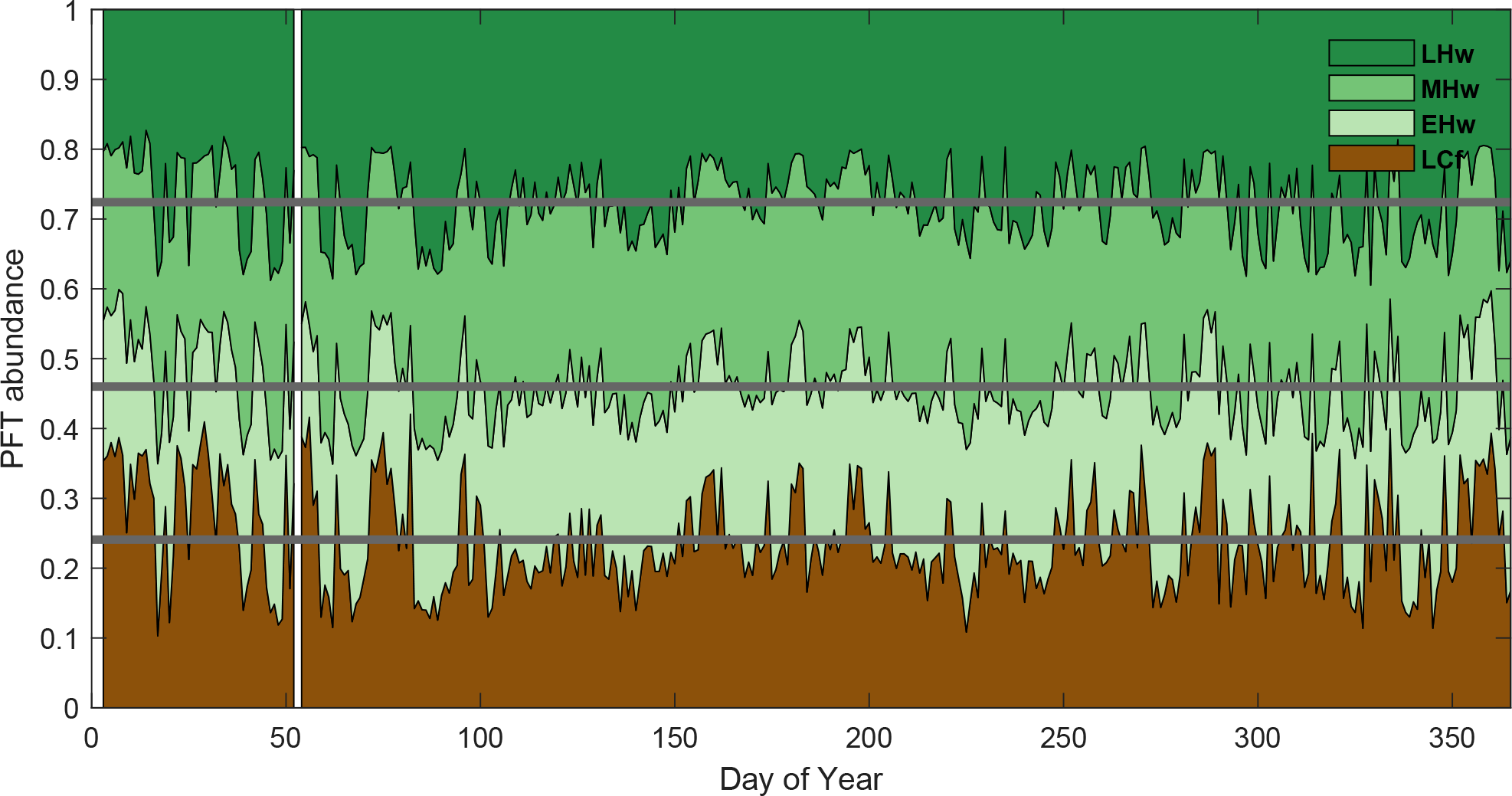
Forest composition as observed by the fluxtower for the year 2016, based on flux footprint predictions. Gray lines mark the annual averages of plant functional type (PFT) abundance that were used to build a representative fluxtower plot representation in the ecosystem demography model.

**Fig. S10:**
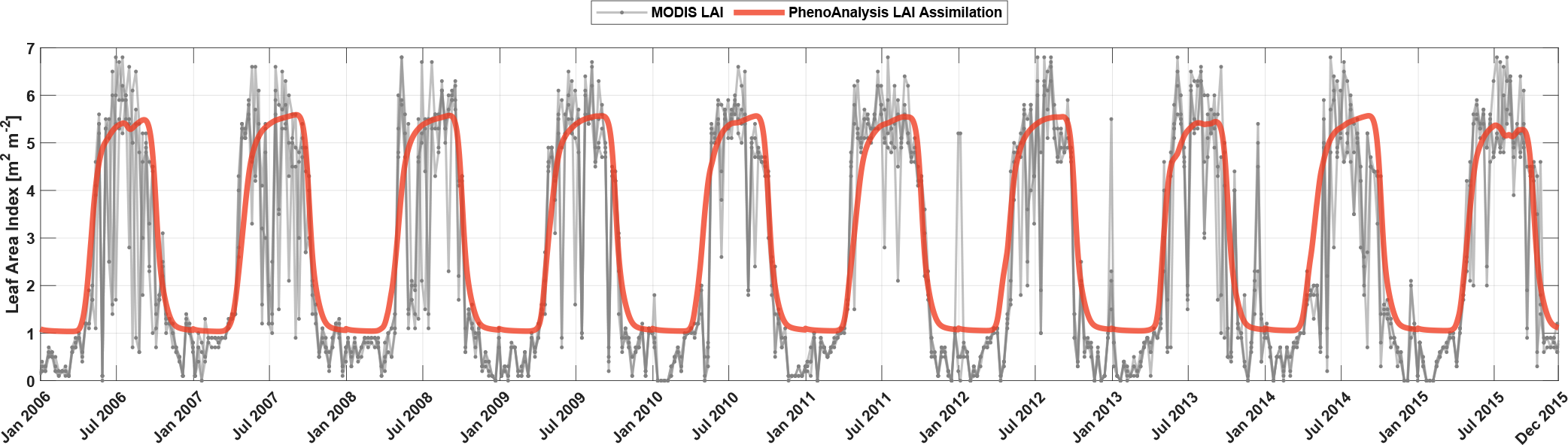
Moderate Resolution Imaging Spectroradiometer (MODIS) leaf area index (LAI) time series for the years 2006–2015 and corresponding modeled LAI using the PhenoAnalysis data assimilation model.

**Fig. S11:**
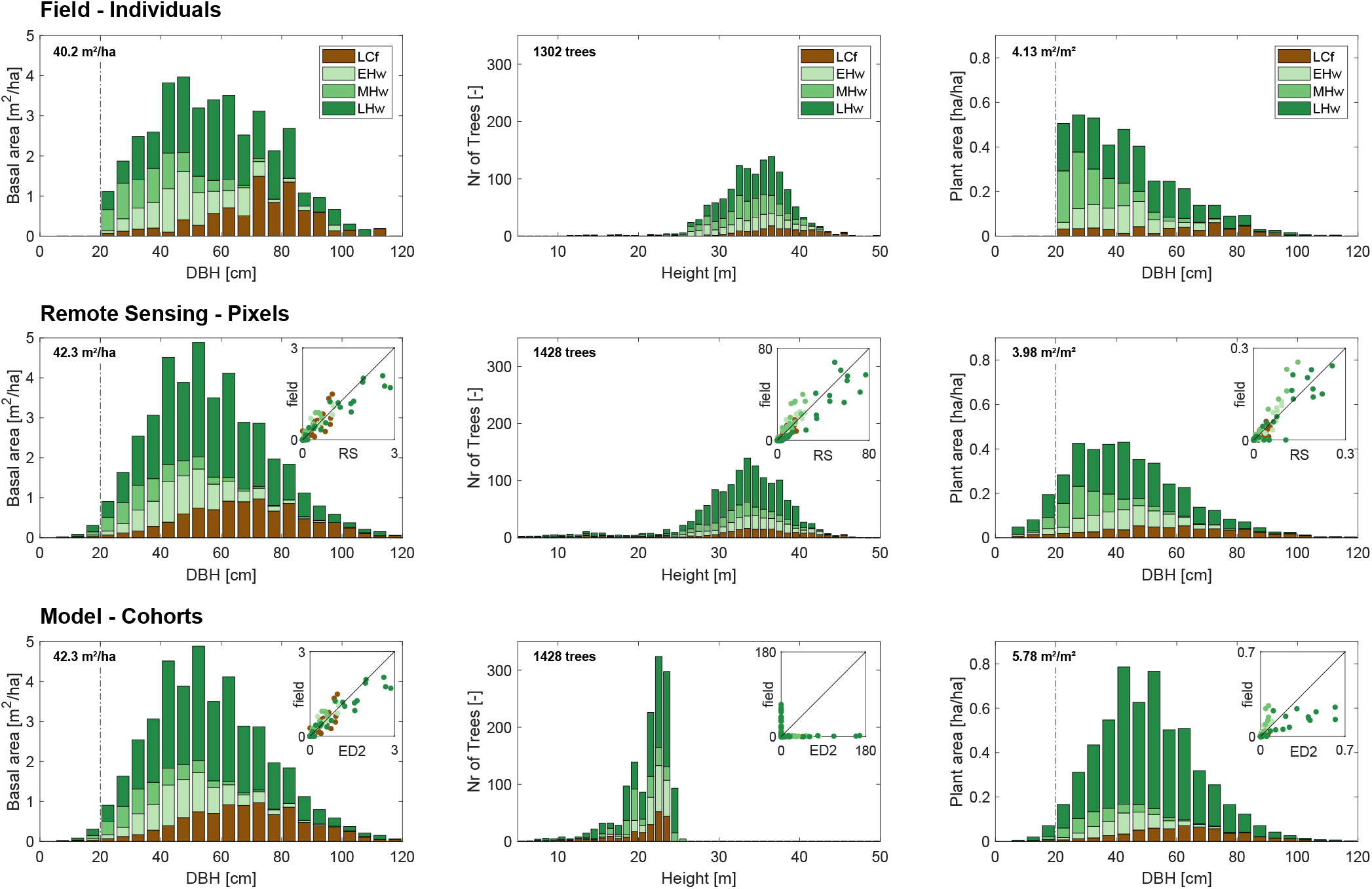
Forest structure and composition measured in the field based on individual tress (trees *>* 20 cm DBH, marked by dashed line), derived from remote sensing data at pixel-level, and represented in the model as cohorts. Note that basal area is a direct input to the model, whereas height and leaf area index are internally derived following PFT-based model allometries. Structure and composition are shown as basal area and plant area per DBH size class and tree count per canopy height class for late-successional conifers (LCf) and early, mid and late-successional hardwoods (EHw, MHw, LHw).

**Fig. S12:**
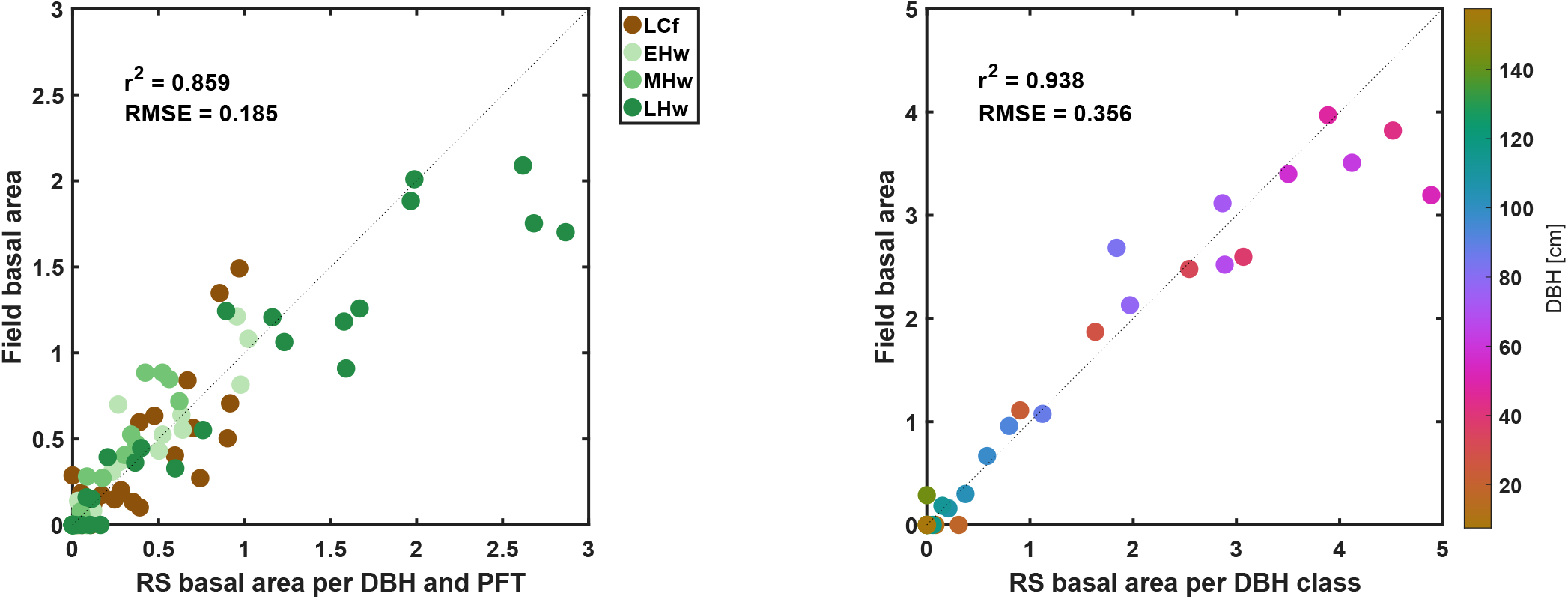
Basal area derived from remote sensing (RS) compared to values measured in the field for late-successional conifers (LCF), early- (EHw), mid- (MHw) and late-successional (LHw) hardwoods (left panel) and in total (right panel), where each dot represents a diameter class (DBH classes 20-150 cm in 10 cm steps).

**Fig. S13:**
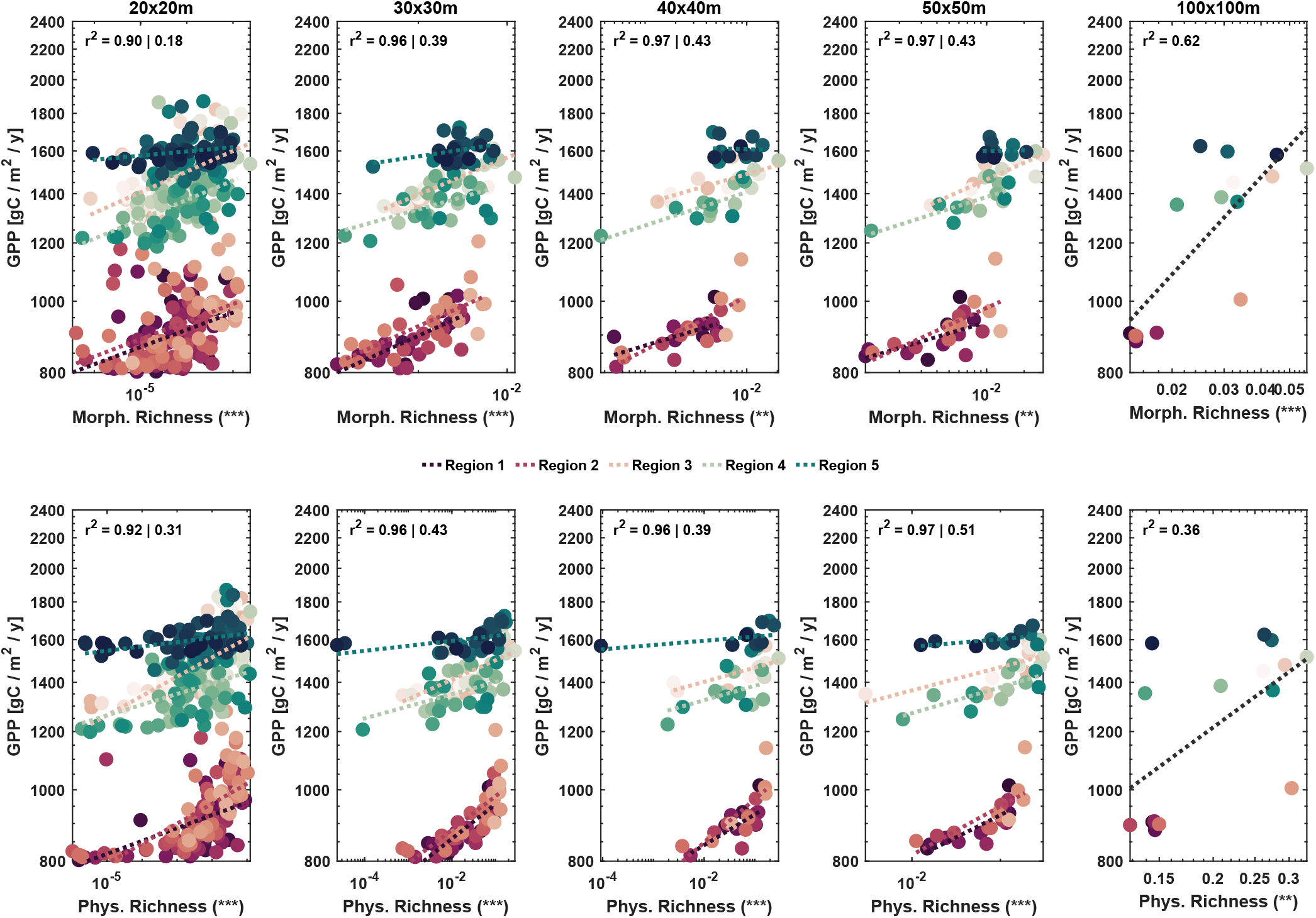
Relationship between morphological (top) and physiological (bottom) richness and productivity as average annual gross primary productivity (GPP), simulated across five regions over the course of a decade with vegetation dynamics and analyzed at five different spatial extents. Underlying functional traits are leaf area index, basal area and foliage height diversity (morphological) and VC*_max_* and specific leaf area (physiological) at 10 m spatial grain (model patch level averages). Significance (*p*) and *r*^2^ *left* are given based on a general linear model controlling for soil/region, while *r*^2^ *right* shows the average of five individual linear models fitted to each region. At 1-ha scale, only one linear model was fitted to the 15 sites.

**Fig. S14:**
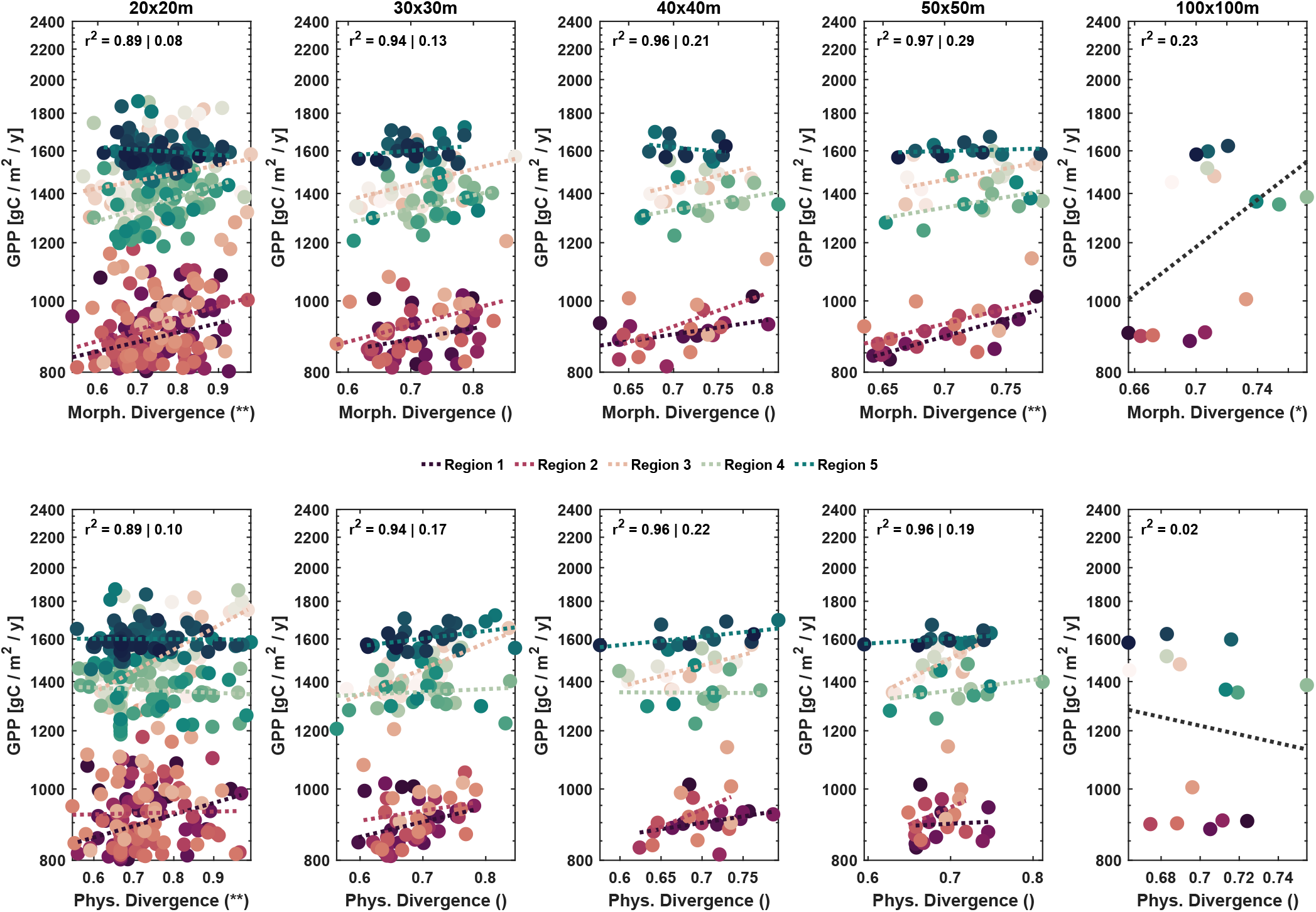
Relationship between morphological (top) and physiological (bottom) divergence and productivity as average annual gross primary productivity (GPP), simulated across five regions over the course of a decade with vegetation dynamics and analyzed at five different spatial extents. Underlying functional traits are leaf area index, basal area and foliage height diversity (morphological) and VC*_max_* and specific leaf area (physiological) at 10 m spatial grain (model patch level averages). Significance (*p*) and *r*^2^ *left* are given based on a general linear model controlling for soil/region, while *r*^2^ *right* shows the average of five individual linear models fitted to each region. At 1-ha scale, only one linear model was fitted to the 15 sites.

**Fig. S15:**
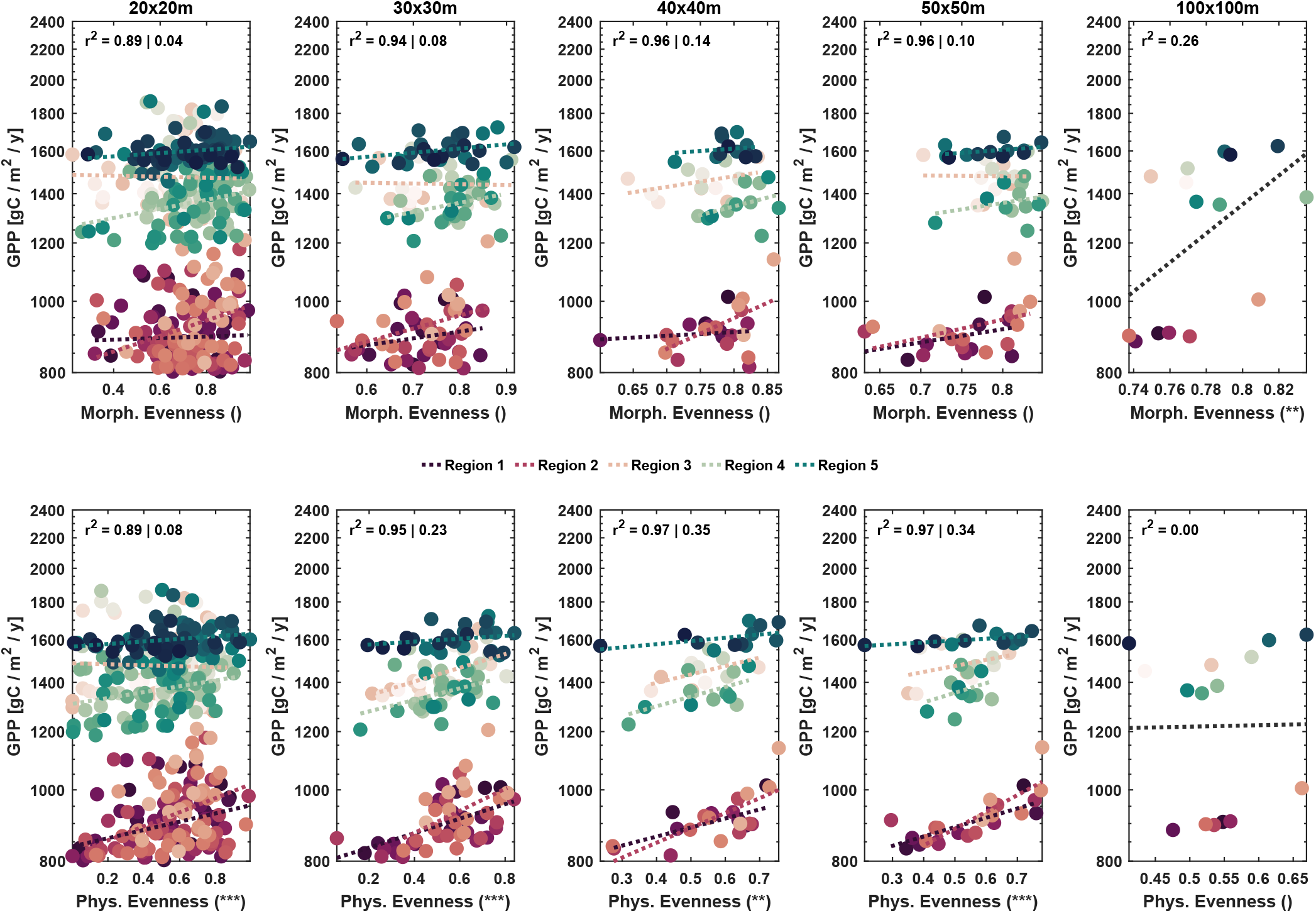
Relationship between morphological (top) and physiological (bottom) evenness and productivity as average annual gross primary productivity (GPP), simulated across five regions over the course of a decade with vegetation dynamics and analyzed at five different spatial extents. Underlying functional traits are leaf area index, basal area and foliage height diversity (morphological) and VC*_max_* and specific leaf area (physiological) at 10 m spatial grain (model patch level averages). Significance (*p*) and *r*^2^ *left* are given based on a general linear model controlling for soil/region, while *r*^2^ *right* shows the average of five individual linear models fitted to each region. At 1-ha scale, only one linear model was fitted to the 15 sites.

**Fig. S16:**
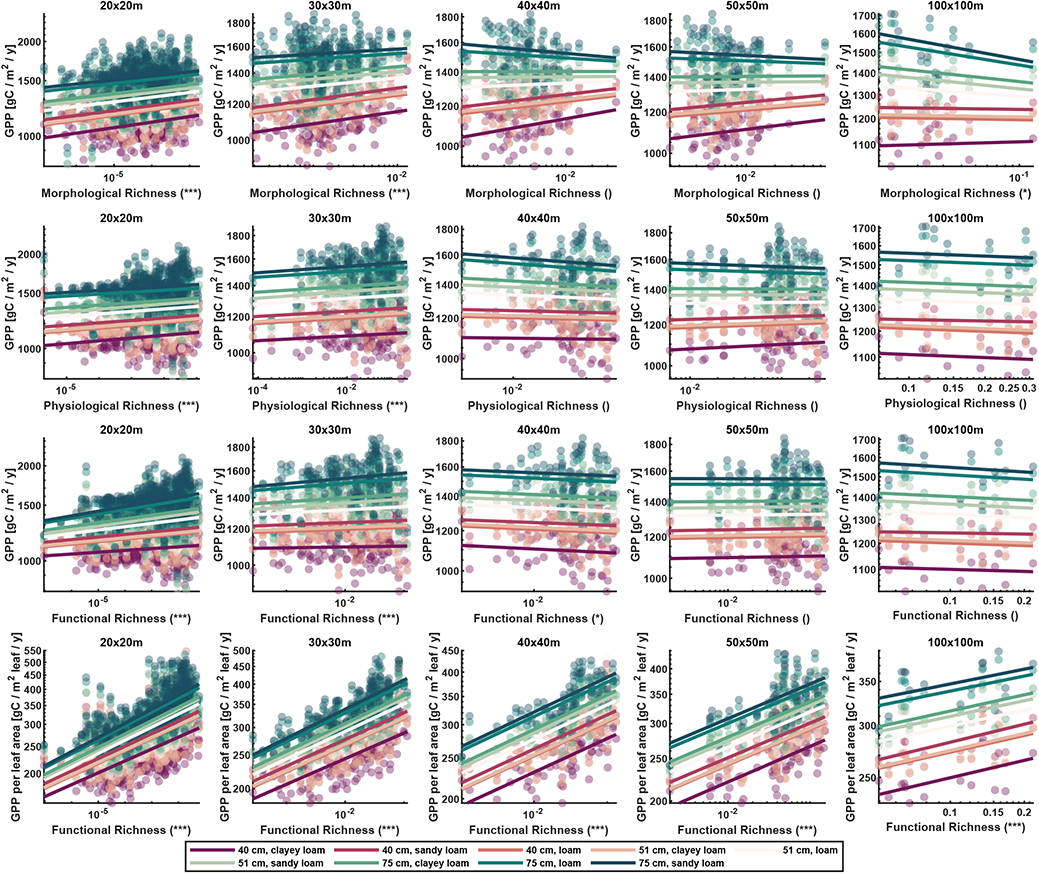
Relationships between morphological, physiological and functional richness and average gross primary productivity (GPP) and GPP normalized by leaf area (bottom row) among nine different soils (colors from red to green) and five different spatial scales (20-100 m). Significance of richness on GPP as determined by a general linear model in log-log space with soil and richness as fixed effects is indicated as *p <* 0.001 (***), *p <* 0.01 (**), *p <* 0.05 (*) and *p >*= 0.05 ().

**Fig. S17:**
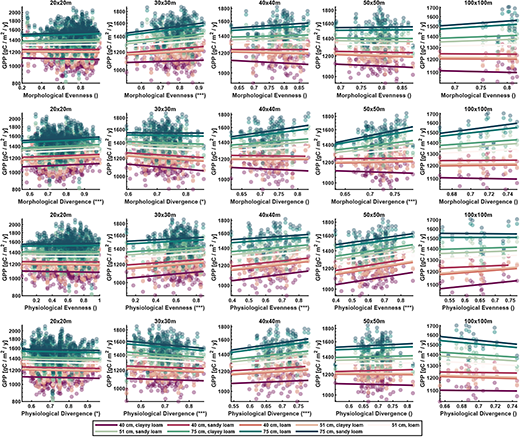
Relationships between morphological and physiological evenness and divergence and average gross primary productivity (GPP) among nine different soils (colors from red to green) and five different spatial scales (20-100 m). Significance of evenness and divergence on GPP as determined by a general linear model in log-log space with soil and evenness or divergence as fixed effects is indicated as *p <* 0.001 (***), *p <* 0.01 (**), *p <* 0.05 (*) and *p >*= 0.05 ().

**Fig. S18:**
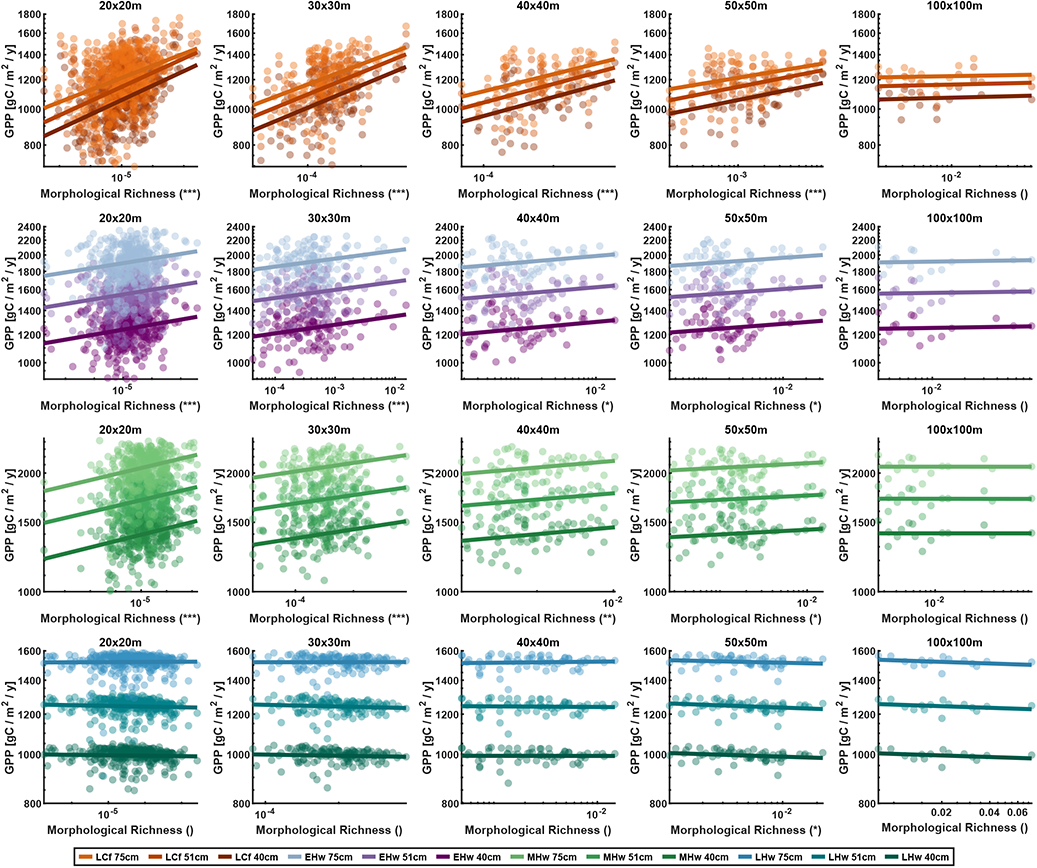
Relationships between morphological richness and average gross primary productivity (GPP) simulated for four monocultures among three different soils (color shading) and five different spatial scales (20-100 m). Monocultures are late-successional conifers (LCf, orange, first row), early-successional hardwoods (EHw, purple, second row), mid-successional hardwoods (MHw, green, third row) and late-successional hardwoods (LHw, blue, fourth row). Significance of morphological richness on GPP as determined by a general linear model in log-log space with soil and richness as fixed effects is indicated as *p <* 0.001 (***), *p <* 0.01 (**), *p <* 0.05 (*) and *p >*= 0.05 ().

**Fig. S19:**
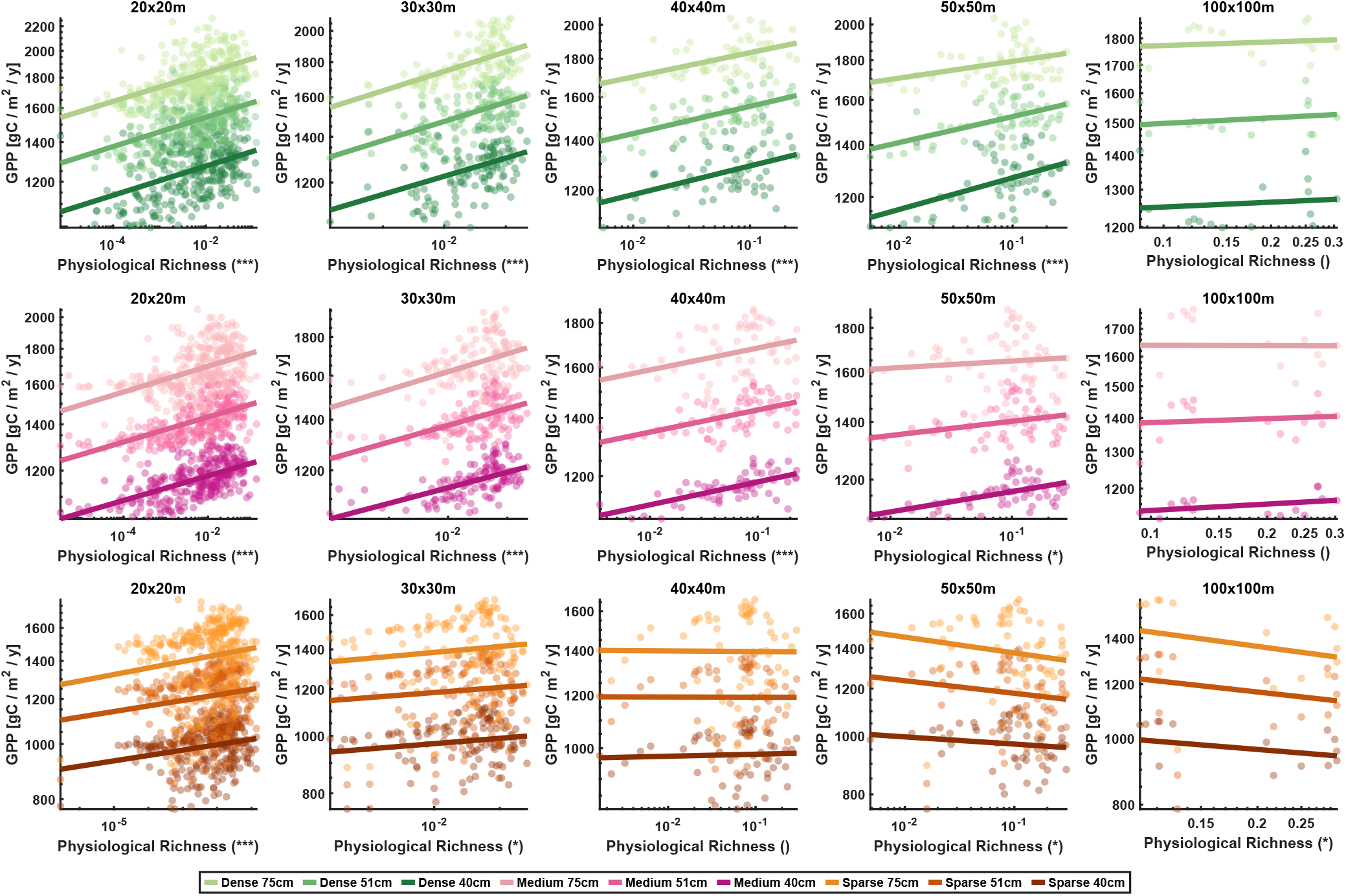
Relationships between physiological richness and average gross primary productivity (GPP) simulated for three mono-structures among three different soils (color shading) and five different spatial scales (20-100 m). Mono-structures are dense (83 m^2^ ha*^−^*^1^, green, first row), medium (39 m^2^ ha*^−^*^1^, pink, second row), and sparse 19 m^2^ ha*^−^*^1^), orange, third row). Significance of physiological richness on GPP as determined by a general linear model in log-log space with soil and richness as fixed effects is indicated as *p <* 0.001 (***), *p <* 0.01 (**), *p <* 0.05 (*) and *p >*= 0.05 ().

**Tab. S1:** Coefficients *a* and *b* of the exponential function and standard deviation *std*, skewness *skew* and kurtosis *kurt* describing the canopy height to dbh allometry for the plant functional types late conifers (LCf), early hardwoods (EHw), mid hardwoods (MHw) and late hardwoods (LCf).

|  | LCf | EHw | MHw | LHw |
| --- | --- | --- | --- | --- |
| $a$ | 11.41 | 17.31 | 16.63 | 11.32 |
| $b$ | 0.466 | 0.280 | 0.225 | 0.398 |
| $std$ | 18.26 | 10.70 | 9.752 | 16.13 |
| $skew$ | -0.475 | 1.226 | 1.106 | 0.443 |
| $kurt$ | 3.705 | 6.973 | 4.332 | 2.959 |

**Tab. S2:** Model: GPP *∼* soil*MRic; morphological richness (MRic) across five different regions / soils (soil), vegetation dynamics on; \**p <* 0.05, \*\**p <* 0.01, \*\*\**p <* 0.001.

| Term | 20 x 20 m |  | 30 x 30 m |  | 40 x 40 m |  | 50 x 50 m |  |
| --- | --- | --- | --- | --- | --- | --- | --- | --- |
| | $F$ | $p$ | $F$ | $p$ | $F$ | $p$ | $F$ | $p$ |
| soil | 21.1 | <0.001*** | 7.63 | <0.001*** | 5.14 | 0.002** | 6.25 | <0.001*** |
| MRic | 18.3 | <0.001*** | 27.8 | <0.001*** | 9.14 | 0.004** | 9.70 | 0.003** |
| soil:MRic | 2.30 | 0.058 | 1.58 | 0.183 | 1.96 | 0.114 | 0.93 | 0.454 |

**Tab. S3:** Model: GPP *∼* soil*PRic; physiological richness (PRic) across five different regions / soils (soil), vegetation dynamics on; \**p <* 0.05, \*\**p <* 0.01, \*\*\**p <* 0.001.

| Term | 20 x 20 m |  | 30 x 30 m |  | 40 x 40 m |  | 50 x 50 m |  |
| --- | --- | --- | --- | --- | --- | --- | --- | --- |
| | $F$ | $p$ | $F$ | $p$ | $F$ | $p$ | $F$ | $p$ |
| soil | 178 | <0.001*** | 88.1 | <0.001*** | 46.5 | <0.001*** | 42.2 | <0.001*** |
| PRic | 46.3 | <0.001*** | 47.4 | <0.001*** | 10.0 | 0.003** | 21.4 | <0.001*** |
| soil:PRic | 7.93 | <0.001*** | 11.2 | <0.001*** | 5.98 | <0.001*** | 3.74 | 0.010** |

**Tab. S4:** Model: GPP *∼* soil*MDiv; morphological divergence (MDiv) across five different regions / soils (soil), vegetation dynamics on; \**p <* 0.05, \*\**p <* 0.01, \*\*\**p <* 0.001.

| Term | 20 x 20 m |  | 30 x 30 m |  | 40 x 40 m |  | 50 x 50 m |  |
| --- | --- | --- | --- | --- | --- | --- | --- | --- |
| | $F$ | $p$ | $F$ | $p$ | $F$ | $p$ | $F$ | $p$ |
| soil | 17.5 | <0.001*** | 4.54 | 0.002** | 3.96 | 0.007** | 4.41 | 0.004** |
| MDiv | 6.76 | 0.010** | 3.34 | 0.070 | 3.41 | 0.071 | 16.0 | 0.004** |
| soil:MDiv | 2.13 | 0.077 | 0.41 | 0.799 | 1.45 | 0.230 | 1.11 | 0.363 |

**Tab. S5:** Model: GPP *∼* soil*PDiv; physiological divergence (PDiv) across five different regions / soils (soil), vegetation dynamics on; \**p <* 0.05, \*\**p <* 0.01, \*\*\**p <* 0.001.

| Term | 20 x 20 m |  | 30 x 30 m |  | 40 x 40 m |  | 50 x 50 m |  |
| --- | --- | --- | --- | --- | --- | --- | --- | --- |
|  | *F* | *p* | *F* | *p* | *F* | *p* | *F* | *p* |
| soil | 21.5 | <0.001*** | 5.11 | <0.001*** | 4.12 | 0.006** | 2.19 | 0.084 |
| PDiv | 9.49 | 0.002** | 3.84 | 0.052 | 1.51 | 0.226 | 0.09 | 0.085 |
| soil:PDiv | 9.25 | <0.001*** | 1.83 | 0.128 | 1.42 | 0.240 | 1.46 | 0.227 |

**Tab. S6:** Model: GPP *∼* soil*MEve; morphological evenness (MEve) across five different regions / soils (soil), vegetation dynamics on; \**p <* 0.05, \*\**p <* 0.01, \*\*\**p <* 0.001.

| Term | 20 x 20 m |  | 30 x 30 m |  | 40 x 40 m |  | 50 x 50 m |  |
| --- | --- | --- | --- | --- | --- | --- | --- | --- |
|  | *F* | *p* | *F* | *p* | *F* | *p* | *F* | *p* |
| soil | 38.7 | <0.001*** | 10.1 | <0.001*** | 3.36 | 0.016* | 2.22 | 0.081 |
| MEve | 0.15 | 0.695 | 2.02 | 0.158 | 0.22 | 0.644 | 2.20 | 0.144 |
| soil:MEve | 2.24 | 0.065 | 1.10 | 0.360 | 1.49 | 0.220 | 0.35 | 0.846 |

**Tab. S7:**
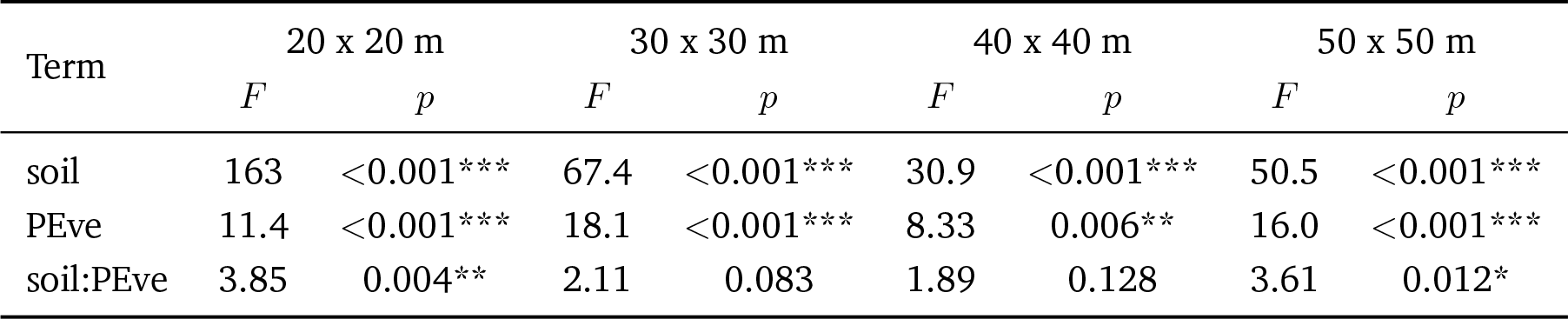
Model: GPP *∼* soil*PEve; physiological evenness (PEve) across five different regions / soils (soil), vegetation dynamics on; \**p <* 0.05, \*\**p <* 0.01, \*\*\**p <* 0.001.

**Tab. S8:**
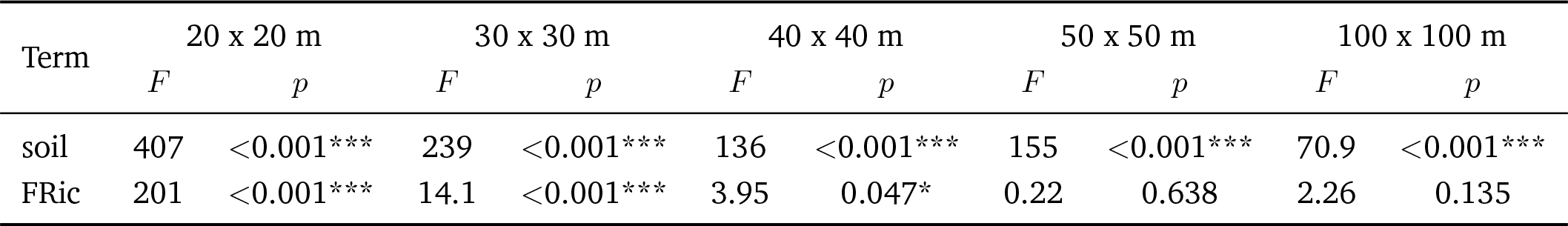
Model: GPP *∼* soil + FRic; functional richness (FRic) across nine mono-soils (soil), vegetation dynamics off; \**p <* 0.05, \*\**p <* 0.01, \*\*\**p <* 0.001.

**Tab. S9:** Model: GPP*_leaf_ ∼* soil + FRic; functional richness (FRic) across nine mono-soils (soil), vegetation dynamics off; \**p <* 0.05, \*\**p <* 0.01, \*\*\**p <* 0.001.

| Term | 20 x 20 m |  | 30 x 30 m |  | 40 x 40 m |  | 50 x 50 m |  | 100 x 100 m |  |
| --- | --- | --- | --- | --- | --- | --- | --- | --- | --- | --- |
|  | *F* | *p* | *F* | *p* | *F* | *p* | *F* | *p* | *F* | *p* |
| soil | 219 | <0.001*** | 156 | <0.001*** | 94.5 | <0.001*** | 86.6 | <0.001*** | 52.9 | <0.001*** |
| FRic | 2433 | <0.001*** | 1409 | <0.001*** | 522 | <0.001*** | 375 | <0.001*** | 56.4 | <0.001*** |

**Tab. S10:** Model: GPP *∼* PFT + soil + PFT:soil + MRic; morphological richness (MRic) across four monocultures (PFT) and three mono-soils (soil), vegetation dynamics off; \**p <* 0.05, \*\**p <* 0.01, \*\*\**p <* 0.001.

| Term | 20 x 20 m |  | 30 x 30 m |  | 40 x 40 m |  | 50 x 50 m |  | 100 x 100 m |  |
| --- | --- | --- | --- | --- | --- | --- | --- | --- | --- | --- |
|  | *F* | *p* | *F* | *p* | *F* | *p* | *F* | *p* | *F* | *p* |
| PFT | 586 | <0.001*** | 325 | <0.001*** | 157 | <0.001*** | 209 | <0.001*** | 92.5 | <0.001*** |
| soil | 94.4 | <0.001*** | 52.5 | <0.001*** | 28.0 | <0.001*** | 34.0 | <0.001*** | 15.7 | <0.001*** |
| PFT:soil | 232 | <0.001*** | 121 | <0.001*** | 59.1 | <0.001*** | 83.4 | <0.001*** | 39.1 | <0.001*** |
| MRic | 279 | <0.001*** | 163 | <0.001*** | 54.1 | <0.001*** | 32.1 | 0.016* | 0.09 | 0.762 |

**Tab. S11:** Model: GPP *∼* CST + soil + CST:soil + PRic; physiological richness (PRic) across three mono-structures (CST) and three mono-soils (soil), vegetation dynamics off; \**p <* 0.05, \*\**p <* 0.01, \*\*\**p <* 0.001.

| Term | 20 x 20 m |  | 30 x 30 m |  | 40 x 40 m |  | 50 x 50 m |  | 100 x 100 m |  |
| --- | --- | --- | --- | --- | --- | --- | --- | --- | --- | --- |
|  | *F* | *p* | *F* | *p* | *F* | *p* | *F* | *p* | *F* | *p* |
| CST | 416 | <0.001*** | 213 | <0.001*** | 116 | <0.001*** | 104 | <0.001*** | 35.3 | <0.001*** |
| soil | 1734 | <0.001*** | 753 | <0.001*** | 391 | <0.001*** | 336 | <0.001*** | 110 | <0.001*** |
| CST:soil | 17.9 | <0.001*** | 7.91 | <0.001*** | 4.41 | 0.002** | 3.65 | 0.006** | 1.20 | 0.313 |
| PRic | 414 | <0.001*** | 134 | <0.001*** | 30.3 | <0.001*** | 5.84 | 0.016* | 1.20 | 0.275 |

### Supplementary Note 1: Remote Sensing Data

Full-waveform airborne laser scanning data based on light detection and ranging (LiDAR) was acquired in the period of 19 June to 25 July 2014 as part of a larger flight campaign covering the whole Kanton Aargau. The scanner system with a rotating mirror (RIEGL LMS-Q680i, scan angle *±*22*^◦^*) was flown under leaf-on conditions at a nominal height of 700 m above ground, resulting in a footprint size of approximately 35 cm and an average point density of 30 pts/m^2^. The LiDAR data was registered to the Swiss national grid CH1903+ with a positional accuracy of *<* 0.07 m in vertical and *<* 0.15 m in horizontal direction. For more details about the acquisition and the quality of the data, see Kükenbrink *et al*. (2017).

Imaging spectroscopy acquisitions were flown on 7 July and 17 July 2016 under clear sky conditions using the APEX imaging spectrometer (Schaepman *et al*., 2015). The study area was covered with one flight line on each of the acquisition dates. The average flight altitude was 4500 m a.s.l. resulting in an average ground pixel size of 2*×*2-m. APEX measured at-sensor radiances in 284 spectral bands ranging from 399 nm to 2425 nm. APEX data were processed to hemispherical-conical reflectance factors (HCRF) in the APEX Processing and Archiving Facility (Hueni *et al*., 2009). Processing started with the raw instrument data, which was split into image, dark current and housekeeping data, thus forming level 0. Level 1 (L1) calibrated radiances were obtained by inverting the instrument model, applying coefficients established during calibration and characterization at the APEX Calibration Home Base (Hueni *et al*., 2013). The position and orientation of each pixel in 3-D space was based on automatic geocoding in PARGE v3.3 (Schläpfer & Richter, 2002), using the swissALTI3D digital terrain model and automatic image co-registration to swissimage ortho-photos. L1 data were converted to HCRF by employing ATCOR4 v7.1 in the smile aware mode (Schaepman *et al*., 2015). Finally, HCRF surface reflectance values were corrected by a vicarious calibration based on 27 ground reflectance targets with various brightness levels located within the flight stripes to minimize remaining radiometric differences between overflights. Ground targets were measured using a field spectroradiometer (ASD Fieldspec Pro 4).

Remotely sensed physiological and morphological trait data were used from Schneider *et al*. (2017). Specifically, we used functional trait maps of the Laegern forest with relative content of chlorophyll, carotenoids, and leaf water, as well as plant area index, canopy height, and foliage height diversity with a resolution of 2*×*2-m pixel size. We also used the 3D plant area index voxel grid with 2*×*2*×*2-m resolution. For detailed information about the retrieval of the functional traits and the 3D voxel grid we refer to Schneider *et al*. (2014, 2017).

### Supplementary Note 2: Random Forest PFT Classification

To represent forest composition in the model, we group the species present at our site into corresponding ED2 plant functional types used by Medvigy *et al*. (2009) and Antonarakis *et al*. (2014), namely: late-successional conifers (LCf), and early (EHw), mid (MHw), and late-successional hardwoods (LHw). At the Laegern forest, the two late-successional coniferous needle species are Norway spruce (*Picea abies*) and silver fir (*Abies alba*). Furthermore, we grouped European ash (*Fraxinus excelsior*) and common whitebeam (*Sorbus aria*) as early-successional hardwoods, sycamore maple (*Acer pseudoplatanus*), field maple (*Acer campestre*), sessile oak (*Quercus petraea*), largeleaf linden (*Tilia platyphyllos*) and wych elm (*Ulmus glabra*) as mid-successional hardwoods, and European beach (*Fagus sylvatica*), Norway maple (*Acer platanoides*) and European hornbeam (*Carpinus betulus*) as late-successional hardwoods. There were no early- and mid-successional coniferous species present at our site.

We calculated a set of 81 possible input features based on six functional traits, corresponding spatially filtered traits, airborne imaging spectroscopy and laser scanning data to perform a feature selection. The data are described in Schneider *et al*. (2017) and Supplementary Note 1. Based on feature importance, we discarded features with low importance that reduced the overall accuracy of the random forest model. We received the best results with 27 relevant features. We used the six morphological and physiological forest traits at 2*×*2-m spatial resolution as well as the corresponding filtered traits using a spatial median filter with a window size of 7*×*7 pixels. We performed a principal component analysis (PCA) for forested pixels on the 284 surface reflectance bands and selected the PCA bands 1 to 9, 13 and 18. Additionally, we applied a continuum removal (CR) on the 284 surface reflectance bands followed by a PCA and selected the CR-PCA bands 1, 2, 4 and 10. This proved to be the best set of input features for the random forest classification of plant functional types.

We used the field inventory data of 1307 dominant and co-dominant trees and the corresponding crown map to derive a raster-based training dataset at 2*×*2-m spatial resolution (Morsdorf *et al*., 2020; Guilĺen-Escribà *et al*., 2021). In the original dataset, there were 1449 pixels for late conifers, 1960 pixels for early hardwoods, 1916 pixels for mid hardwoods and 5262 pixels for late hardwoods. We then used prior information from 73 forest stand polygons covering the Laegern forest to build specific training datasets for each polygon. Stand polygons of Kanton Aargau and Zurich include forest stand information on development stage, the percentage coverage of the 6 most dominant species, and the percentage coverage of deciduous broadleaf and coniferous needle trees. The data from Kanton Aargau was provided by Aargauisches Geografisches Informationssystem (AGIS), Departement Bau, Verkehr und Umwelt, Abteilung Wald (last updated on 27 February 2015). The data from Kanton Zurich was provided by Geographisches Informationssystem (GIS-ZH), Amt für Landschaft und Natur, Abteilung Wald (last updated on 16 September 2015).

We adapted the composition of the four plant functional types in the training datasets based on the percentage coverage of the most dominant species according to the stand polygon data. We trained, tested and ran a random forest classification for each stand polygon of the forest with its specific training dataset. We used 70% of the data for training and 30% for testing and validation. We built 1000 trees with a minimum leaf size of 5. We used the composition of groups in the training dataset as prior information for the random forest classification. We then used all of the data to train the model for predicting the plant functional type for each pixel. We reached an overall accuracy of 74% and a kappa of 61% for the classification of plant functional types over the whole forest. For spatially resolved results for each polygon and a comparison between the predicted plant functional types per polygon and the prior information, we refer to Supplementary Figure S3. The final PFT classification map is shown in Supplementary Figure S4.

### Supplementary Note 3: Forest Structure Estimation

The stem segmentation method to estimate stem density is an adapted version of the approach used in Morsdorf *et al*. (2004); Kaartinen *et al*. (2012) and later updated in Wang & Loreau (2016). The algorithm detects local maxima in a smoothed version of the canopy height model (CHM) and uses those positions as starting points for either a clustering of the point cloud (Morsdorf *et al*., 2004) or a watershed segmentation of the CHM, which is what we used for this study. The amount of smoothing was varied between deciduous and coniferous trees using an ALS-based forest class differentiation exploiting differences between echo types and other related metrics. Furthermore, a tree height-crown size heuristic (i.e. larger trees have larger crowns) was used to apply differently sized smoothing kernels for trees of different size. The method had an individual-tree detection rate of 79% overall, 83% for trees *>* 35 m and 76% for trees *<* 35 m. The canopy height map and segmented stems with assigned DBH values are shown in Supplementary Figure S5, and are comparable to the reference data in Supplementary Figure S6. We then related pixel counts to stem counts per site based on 1-m height classes from 4 to 55 m canopy height. This allowed us to estimate the fractional stem count for each pixel. For comparison with the field data, we calculated basal area *ba* (m^2^ ha*^−^*^1^) as follows:

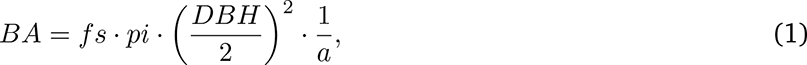

where *fs* is the fractional stem count, *DBH* is the diameter at breast height (cm), and *a* is the total area of the plot (m^2^). Basal area *BA* was calculated for each 2 m pixel and then aggregated to 10 m by summing the pixel values of each plant functional type.

### Supplementary Note 4: Gap filling of meteorological variables

For temperature, wind direction, average wind speed, maximum wind speed and humidity, we used the measurements from the meteorological station at the ridge of the Laegern to fill the gaps in the time series of the fluxtower. We built a linear model based on monthly mean values to fit the values measured on the ridge to the ones measured at the fluxtower. Then we replaced the gaps in the time series by the fitted values from the meteorological station at the ridge. The same procedure was applied for precipitation with data from Kloten, being the closest meteorological station with precipitation measurements. Global radiation did not show a consistent difference between the two measurement stations. Therefore, we used a 5 days window before and after the gap to build the linear model instead of monthly averages. For net radiation, there were no measurements available from nearby stations. We filled the gaps by copying the corresponding period from either before or after the gap. We used global radiation data to build a linear model between the gap and the periods before and after to find out which one was more similar, and to establish a correction factor. We then copied the period from the net radiation data accordingly and applied the correction factor. Atmospheric pressure is a more continuous variable than the other meteorological drivers. Applying the same procedure might thus lead to unrealistic, abrupt changes at the beginning and end of the gap. Therefore, we filled the gaps by copying the values from the station at the Laegern ridge and applying a linear scaling to connect to the values before and after the gap. An example of the gap-filled meteorological data at 10 minutes temporal resolution is shown in Figure S7.

